# Conservation and divergence of canonical and non-canonical imprinting in murids

**DOI:** 10.1101/2022.04.21.488764

**Authors:** Julien Richard Albert, Toshihiro Kobayashi, Azusa Inoue, Ana Monteagudo-Sánchez, Soichiro Kumamoto, Tomoya Takashima, Asuka Miura, Mami Oikawa, Fumihito Miura, Shuji Takada, Masumi Hirabayashi, Keegan Korthauer, Kazuki Kurimoto, Maxim Greenberg, Matthew Lorincz, Hisato Kobayashi

## Abstract

**Background:** Genomic imprinting affects gene expression in a parent-of-origin manner and has a profound impact on complex traits including growth and behaviour. While the rat is widely used to model human pathophysiology, few imprinted genes have been identified in this murid. To systematically identify imprinted genes and genomic imprints in the rat, we used low input methods for genome-wide analyses of gene expression and DNA methylation to profile embryonic and extra-embryonic tissues at allele-specific resolution.

**Results:** We identify 14 and 26 imprinted genes in these tissues, respectively, with 10 of these genes imprinted in both tissues. Comparative analyses with mouse revealed that orthologous imprinted gene expression and associated canonical DNA methylation imprints are conserved in the embryo proper of the Muridae family. However, only 3 paternally expressed imprinted genes are conserved in the extra-embryonic tissue of murids, all of which are associated with non-canonical H3K27me3 imprints. The discovery of 8 novel non-canonical imprinted genes unique to the rat is consistent with more rapid evolution of extra-embryonic imprinting. Meta-analysis of novel imprinted genes revealed multiple mechanisms by which species-specific imprinted expression may be established, including H3K27me3 deposition in the oocyte, the birth of ZFP57 binding motifs and the insertion of endogenous retroviral promoters.

**Conclusions:** In summary, we provide a comprehensive list of imprinted loci in the rat, reveal the extent of conservation of imprinted gene expression, and identify potential mechanisms responsible for the evolution of species-specific imprinting.

## INTRODUCTION

The brown Norway rat, an important model for human pathophysiology (1). To facilitate pharmacogenomic studies and the identification of disease-associated variants, efforts have been made to assemble the rat genome (2) and measure the extent of genetic variability between 40 distinct lab strains (3, 4). However, despite the utility of the rat in modeling human disease and recent advances in rat genomics, the mouse has been and continues to be the predominant model species for foundational studies of mammalian genetics. The discovery of genomic imprinting for example, was enabled by nuclear transfer and genetic technologies developed in the mouse (5–8) and numerous follow-up studies of the molecular basis of imprinting have been carried out using mouse models.

This process of genomic imprinting results in monoallelic gene expression in a parent-of-origin manner and is essential for mammalian growth and development. Recent studies in the mouse reveal that oocyte and sperm chromatin is “imprinted” by differential epigenetic modifications that can be maintained in the embryo and adult (9). Canonical imprinted genes are regulated by parent-specific DNA methylation (DNAme) deposited in spermatozoa or oocytes, resulting in differentially methylated regions (DMRs). Imprinted DMRs are maintained on both alleles of the embryo, conferring parent-of-origin monoallelic expression. While roughly 197 imprinted genes have been identified in mouse, 63 of which are also imprinted in human (10), only 13 imprinted genes have been reported in the rat to date (11). Of note, 12 of these genes, including *H19* and *Igf2*, are regulated by DMRs deposited in the gametes, consistent with canonical imprinting. Furthermore, as all 13 rat imprinted genes were characterized based on homology with their mouse or human orthologs, no rat-specific imprinted genes have been reported. Thus, the full extent of genomic imprinting in rats, and the conservation of imprinting between mammals, remains unclear. The lack of a comprehensive list of imprinted genes in rats also hinders the application of comparative genomics to identify novel conserved genomic features that may contribute to imprinting.

Recent studies in the mouse have shown that an alternative mechanism of genomic imprinting, so-called non-canonical imprinting, mediates paternal-specific gene expression in extra-embryonic tissues (12, 13). Non-canonical imprints are distinguished from canonical imprints by the lack of a DNAme imprint and the enrichment in oocytes of histone 3 lysine 27 trimethylation (H3K27me3), a mark deposited by the Polycomb repressive complex PRC2. Such maternal-specific H3K27me3 is replaced by DNAme in extra-embryonic tissues, resulting in a DMR which restricts expression from the maternal allele (12–15). While the molecular basis of this switch remains unknown, recent studies in the mouse implicate GLP and G9A, which deposit H3K9me2 (16), in maintaining maternal DNAme and imprinted gene expression in extraembryonic tissues (17, 18). Of the ∼7 known non-canonical imprinted genes identified in mice, only *Sfmbt2* has been confirmed as an imprinted gene in rat (11, 19). Whether the other non-canonical imprints, such as that controlling expression of the essential growth-factor gene *Gab1* (12, 13, 15, 20), are conserved, remains an open question (11, 21, 22).

Parent-of-origin specific control of gene dosage is hypothesized to be the ultimate driving factor for the evolution of genomic imprinting (23). At the interface between embryo and mother, the placenta is responsible for nutrient transport to the embryo, and gene dosage in this tissue is critical for modulating resource allocation between the mother and embryo (21). Indeed, the placenta shows the highest degree of imprinted gene expression in human and mouse (24), and non-canonical imprinting is restricted to the placenta (12, 21). In an extreme case, in lieu of random X chromosome inactivation, female mice undergo inactivation of the paternal X chromosome in the placenta (25). However, the prevalence of non-canonical imprinting and imprinted X chromosome inactivation in species other than the house mouse (*Mus musculus*) has not been explored.

While the catalogs of human and mouse imprinted genes have been generated over four decades of clinical, cytogenetic and complementation research (10), recent advances in genomics have enabled the comprehensive identification of candidate imprinted genes. By crossing genetically distinct individuals, F1 hybrids are generated with homologous chromosomes that can be differentiated at the genic level *in silico* using known genetic variants. Bioinformatic pipelines, such as our allele-aware tool for processing epigenomic data (MEA) (26) and others such as WASP (27) and Allelome.PRO (28), input data derived from F1 hybrids and generate maternal-and paternal-genome specific profiles. Importantly, parent-of-origin effects, including genomic imprinting, can be delineated from genetic or strain-specific effects by generating F1s from reciprocal crosses (29). Together, allele-specific analysis of genic expression by RNA sequencing (RNAseq) and DNAme levels by whole genome bisulphite sequencing (WGBS) data derived from reciprocal F1 hybrids represent the gold-standard for identifying candidate imprinted genes genome-wide in a given tissue or cell type (30, 31).

In this study, we performed RNAseq and WGBS on reciprocal F1 hybrid rat extra-embryonic and embryonic tissues from early post-implantation conceptuses. Allele-specific profiling of these datasets yielded comprehensive maps of parent-specific gene expression and DNA methylation. In parallel, we generated similar datasets from matching mouse extra-embryonic and embryonic tissues. Comparisons between rat and mouse revealed conserved canonical genomic imprints in the embryo proper but divergent non-canonical imprinting in the extra-embryonic tissue. Detailed inspection of species-specific non-canonical imprinted loci reveal multiple potential mechanisms by which imprinted expression may evolve, including recent insertions of endogenous retroviral promoters that are subject to maternal-specific methylation following fertilization, as well as novel ZFP57 binding motifs that arise through single nucleotide substitutions. Finally, analysis of H3K27me3 in rat oocytes reveals that species-specific deposition of this mark likely explains the divergent non-canonical imprinting status of some genes in extra-embryonic tissues. Altogether, these results provide a comprehensive view of genomic imprinting, including genome-wide maps of parent-of-origin gene expression and parent-specific DNAme levels in the embryo and extra-embryonic tissue, and reveal relatively recent evolutionary divergence of non-canonical imprinting in murids.

## RESULTS

### Measuring imprinted gene expression in the rat

Genetic variation between parental genomes is a prerequisite for the characterization of the transcriptome and methylome with allele-specific resolution, and in turn the systematic identification of putative imprinted genes. To produce embryos from which genomic imprinting in the rat could be assessed, we conducted reciprocal crosses of genetically distinct rat strains BN/NCrlCrlj, WKY/NCrlCrlj and F344/NSlc (hereafter referred to as “B”, “W” and “F”, respectively) for which whole genome sequences are available (3, 4) (**Fig. 1a**). Post-implantation F1 embryos were dissected at Carnegie stage 7, corresponding to embryonic day 8.5 (E8.5) in rat. To agnostically measure embryonic and extra-embryonic gene expression, we performed strand-specific RNAseq on epiblast and ectoplacental cone (EPC) cells, respectively. Subsequently, to identify imprinted gene expression, we analyzed the reciprocal F1 RNAseq datasets using MEA (26), which discriminates transcripts from each allele based on informative parental SNVs as well as INDELs. Expressed autosomal transcripts (RPKM ≥1) with sufficient allele-specific read coverage (allelic RPM ≥0.5 on either allele in at least 6/11 replicates, n=13,165 and 16,642 transcripts for epiblast and EPC samples, respectively) were categorized as paternally or maternally expressed if they showed statistically significant bias in monoallelic expression (Bonferroni-adjusted p-value <0.05, Student’s t-test). Several genes previously reported as imprinted in distantly related mammalian species including mouse, rat, human and cow such as *Igf2* and *Peg10* (32–35) showed paternal allele-specific transcription in both epiblast and EPC, while *H19* showed maternal-specific expression (**Fig. 1b-c & Sup. fig 1a**), validating our RNAseq-based method and parameters for identifying imprinted genes. In total, 19 paternally and 11 maternally expressed imprinted genes were identified in rat epiblast and EPCs using stringent cutoffs defined above. To identify genes that show a bias in parental expression levels (in addition to monoallelic expression), we performed linear modeling of allele-specific data using Limma. An additional 8 paternally and 8 maternally expressed genes were identified using conservative cutoffs (allelic RPM ≥0.5 on either allele in at least 2 replicates per cross in either tissue, n=26,356 transcripts, Benjamini-Hochberg adjusted p-value <0.05, eBayes F-statistic, ≥4-fold change in expression between alleles) and are included in **Fig. 1d** (see **Sup. table 1** for a full list of imprinted genes). While genes identified using Limma show parent-of-origin expression patterns and include the mouse imprinted gene *Slc38a4*, we focused subsequent analyses on genes that met our stringent statistical cutoffs, as defined above.

**Figure 1.**
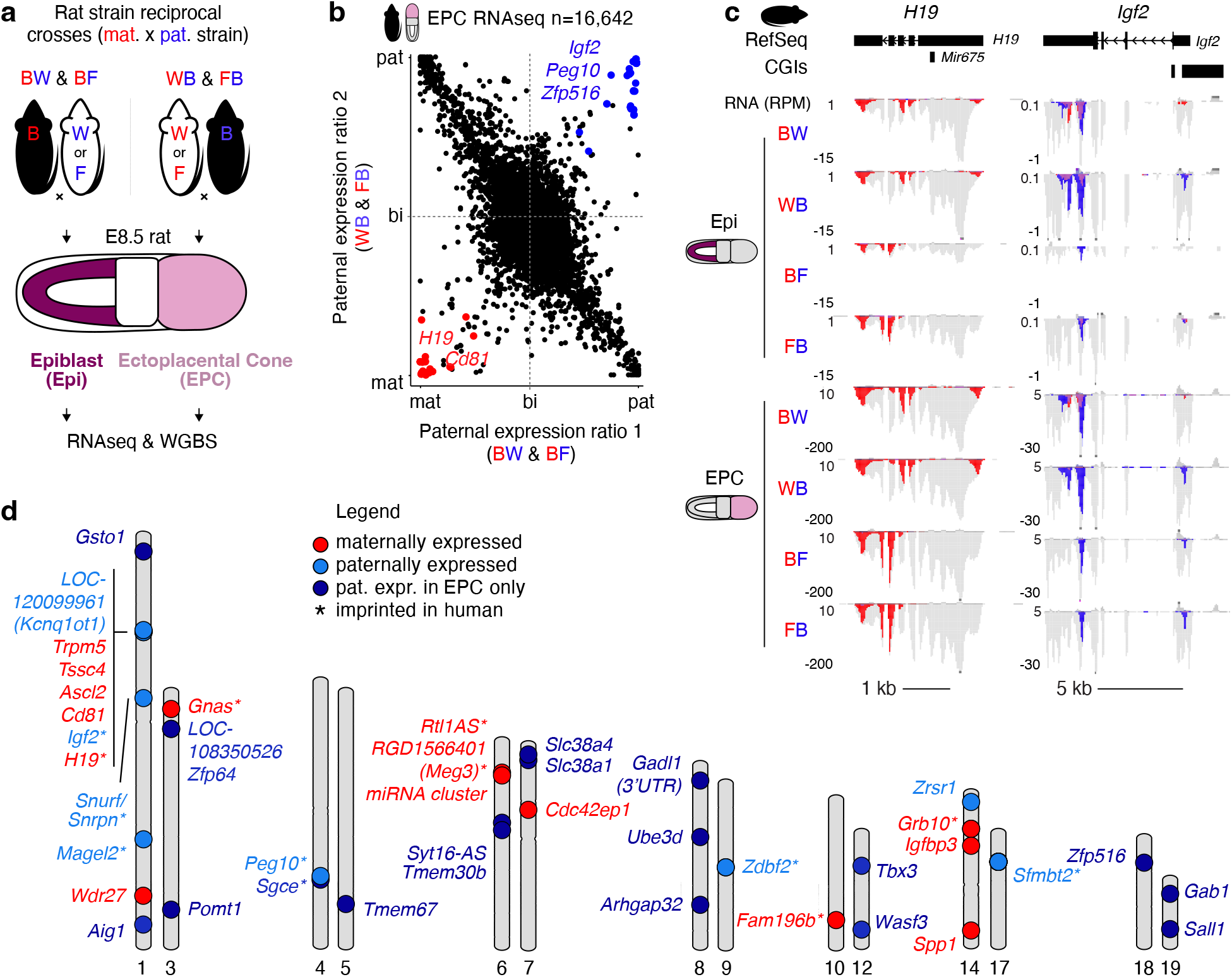
Imprinted gene expression in rat embryonic and extraembryonic cells. **a** Experimental design. Two distinct reciprocal crosses of rat strains (BN/CrlCrlj, “B” and WKY/NCrlCrlj, “W” and F344/NSlc, “F”) were conducted and cells from the E8.5 epiblast (Epi) and the ectoplacental cone (EPC) were collected. RNAseq was performed on all samples (in duplicate or triplicate) and WGBS was performed in duplicate on BW/WB matings. The maternal (red) strain is listed first in cross names. **b** Scatterplot of paternal expression ratios of transcripts in rat EPCs. The paternal expression ratio was averaged over 11 samples generated from BW, BF, WB & FB matings. Expressed transcripts (RPKM ≥1) with sufficient allelic coverage (RPM ≥0.5) in at least 6 samples are shown (n=16,642). Transcripts showing parent-of-origin imprinted gene expression (Student’s T-test, Bonferroni-adjusted p-val <0.05) are coloured red (maternally expressed) or blue (paternally expressed). **c** Rat genome browser screenshots of the maternally expressed imprinted gene H19 and paternally expressed imprinted gene Igf2. The genomic position of known Refseq genes and CpG islands (CGIs) are included. For each cross, duplicates or triplicates were merged and the mean expression level is displayed in reads per million (RPM). Read alignments (grey) are highlighted if they align to the maternal (red) or paternal (blue) alleles. **d** Ideogram karyotype summary of imprinted gene expression identified using a combination of T-test and Limma in rat Epi and EPC cells. Genes that show imprinted gene expression in human are indicated with an asterisk. Genes showing maternal imprinted expression uniquely in EPC cells and normally expressed in adult rat blood (allele-agnostic RPKM ≥1) are not shown (see **Sup. table. 1** for a full list of imprinted genes).

Consistent with previous findings in mouse and human (36), putative rat canonical imprinted genes show parent-specific expression in epiblast and EPCs and are positioned in clusters such as *H19*/*Igf2, Trpm5/Tssc4/Alsc2/Cd81* and *Peg10/Sgce* (**Fig. 1d**). In contrast, putative non-canonical imprinted genes such as *Sfmbt2, Gab1* and *Sall1* show paternal-specific expression exclusively in EPCs. Thus, allele-specific expression analysis of epiblast and EPCs confirmed the classification of known canonical and non-canonical imprinted genes. Furthermore, we identified 8 novel imprinted genes in the rat, including *Zfp516, Slc38a1, Zfp64, Gsto1, Rpl39l, Syt16AS, Gadl1-3’UTR*, and *LOC108350526*. As these genes have not been reported to be imprinted in other mammals, we chose to characterize them in greater detail, as described below.

### Identification of canonical differentially methylated regions in the rat

Having identified known and novel imprinted genes in rat based on allele-specific expression patterns, we next sought to locate candidate regulatory regions responsible for their parent-of-origin transcriptional regulation. Towards this end, we conducted WGBS on the same cell types (see **Sup. table 3** for the complete list of data generated and analyzed in this study) and used MEA to generate parent-of-origin specific methylomes. Subsequently, DSS (37, 38) was employed to identify allele-specific DMRs. Using stringent statistical significance (p-adj <0.001) but lenient DNAme difference (delta >10%) filters, 45,119 DMRs were identified in EPC samples, whereas only 9,165 were identified in epiblasts (**Sup. table 2**). Notably, while the vast majority of DMRs found in EPCs are maternally methylated (n=40,427, 90%), a roughly equal number of paternal and maternal DMRs were identified in epiblast samples (maternally methylated n=4,386, 48%). To distinguish imprints established in the gametes from those that arise following fertilization, we subsequently analyzed our previously generated rat oocyte and sperm WGBS datasets (39). Integrated analysis of DNAme levels revealed a total of 5,876 regions that show dramatic DNAme differences (delta ≥50%) between gametes and the parental alleles of epiblast or EPCs (**Sup. Fig. 1b & Sup. table 2**). While 56% (270/484) of DMRs in epiblasts were apparently established in gametes, 50% (2,749/5,462) of DMRs in EPCs were hypermethylated in both gametes. Importantly, 45 DMRs are shared between all samples (**Sup. Fig. 1c**), a subset of which overlap CpG islands (CGIs) located near known imprinted genes, such as the *Peg3*:CGI-promoter, *Commd1*/*Zrsr1*:CGI-promoter (**Sup. Fig. 1d**), *Grb10*:CGI-promoter, *Impact*:CGI-intragenic, *Mest*:CGI-promoter, *Snrpn*:promoter, *H19*:promoter and *Kcnq1*:intragenic. Notably, DMRs were also identified near the promoters of novel candidate rat-specific imprinted genes such as *Zfp516, Zfp64* and *Syt16-AS* (**Sup. fig. 1e**). These results validate our agnostic approach for discovering DMRs between parental alleles of F1 hybrid rats, and reaffirm that canonical imprinting of the previously described imprinted genes listed above likely arose in a common ancestor over 90 million years ago.

### Evolutionary conservation of imprinting in rat and mouse

We next wished to determine whether any of the candidate novel imprinted genes identified in the rat are imprinted in the mouse, using the same genome-wide approach used for the rat. We generated RNAseq and WGBS data from reciprocal crosses of the well characterized mouse strains C57BL/6N and JF1/Ms, hereafter referred to as “C” and “J”, respectively (**Sup. fig. 2a**). Embryos were again dissected at Carnegie stage 7, corresponding to E7.25 in mouse. Rat embryos at this stage were larger and more elongated relative to mouse (**Sup. fig. 2b-c**), as previously observed (40, 41). Analysis of allele-agnostic levels of transcript expression revealed a high concordance between replicates and reciprocal crosses (Spearman correlation >0.93) in both rat and mouse (**Sup. fig. d-e**), indicating these data are highly reproducible. Using the Ensembl (Biomart) homologous gene annotation and hierarchical clustering of gene expression levels in rat and mouse showed greater variation between epiblast and EPC cell types in each species than inter-species variation of the same cell type (**Sup. fig. 2f**). These observations reveal that epiblast and EPC transcriptional programmes are distinct and strongly conserved between rat and mouse and confirm that rat and mouse samples were indeed collected at roughly matching developmental stages. Importantly, known imprinted genes such as *H19* and *Igf2* showed allele-specific expression in mouse epiblast and EPCs, as expected (**Sup. fig. 2g**). Next, we generated parent-of-origin-specific methylomes of the same reciprocal crosses using MEA and integrated existing oocyte (42) and sperm (43) WGBS data. As expected, the *H19/Igf2* gDMR shows maternal-specific methylation in all datasets in mouse, reflecting canonical imprinting (**Sup. fig. 2g**). Together, these data provide a rich resource to investigate canonical and non-canonical genomic imprinting conservation between two species that diverged only ∼13 million years ago (44).

Integrated analysis of matching expression and DNAme data enabled the unambiguous classification of rat genes into two categories: canonical and non-canonical imprinted genes, the latter category being defined by a lack of a germline DMR and imprinted expression restricted to the paternal allele in extraembryonic cells (12). To measure conservation of imprinted gene expression, as well as allele-specific methylation over nearby DMRs, we compared parent-of-origin gene expression and DNAme levels between rat and mouse homologous genes. 21 known imprinted genes showed imprinted gene expression in both species, including 18 canonical (*Snrpn, Peg10, Igf2, Zrsr1, Sgce, Magel2, Zdbf2, Tssc4, Cd81, Ascl2, Trpm5, Mir675, H19, Grb10, Maged2, Meg3, Rtl1AS* and *Gnas*) (**Fig. 2a**), many of which are adjacent to gametic DMRs in both species (**Fig. 2b**), as well as 3 non-canonical (*Sfmbt2, Sall1* and *Gab1*) genes and nearby EPC-specific DMRs. The conserved imprinting status of *Snrpn, Magel2, Peg10, Sgce, Igf2, Zdbf2, Mir675, H19, Grb10, Meg3, Rtl1AS* and *Gnas* was expected, as they are also imprinted in humans, consistent with an origin of imprinting in an ancient common ancestor (45–47).

**Figure 2.**
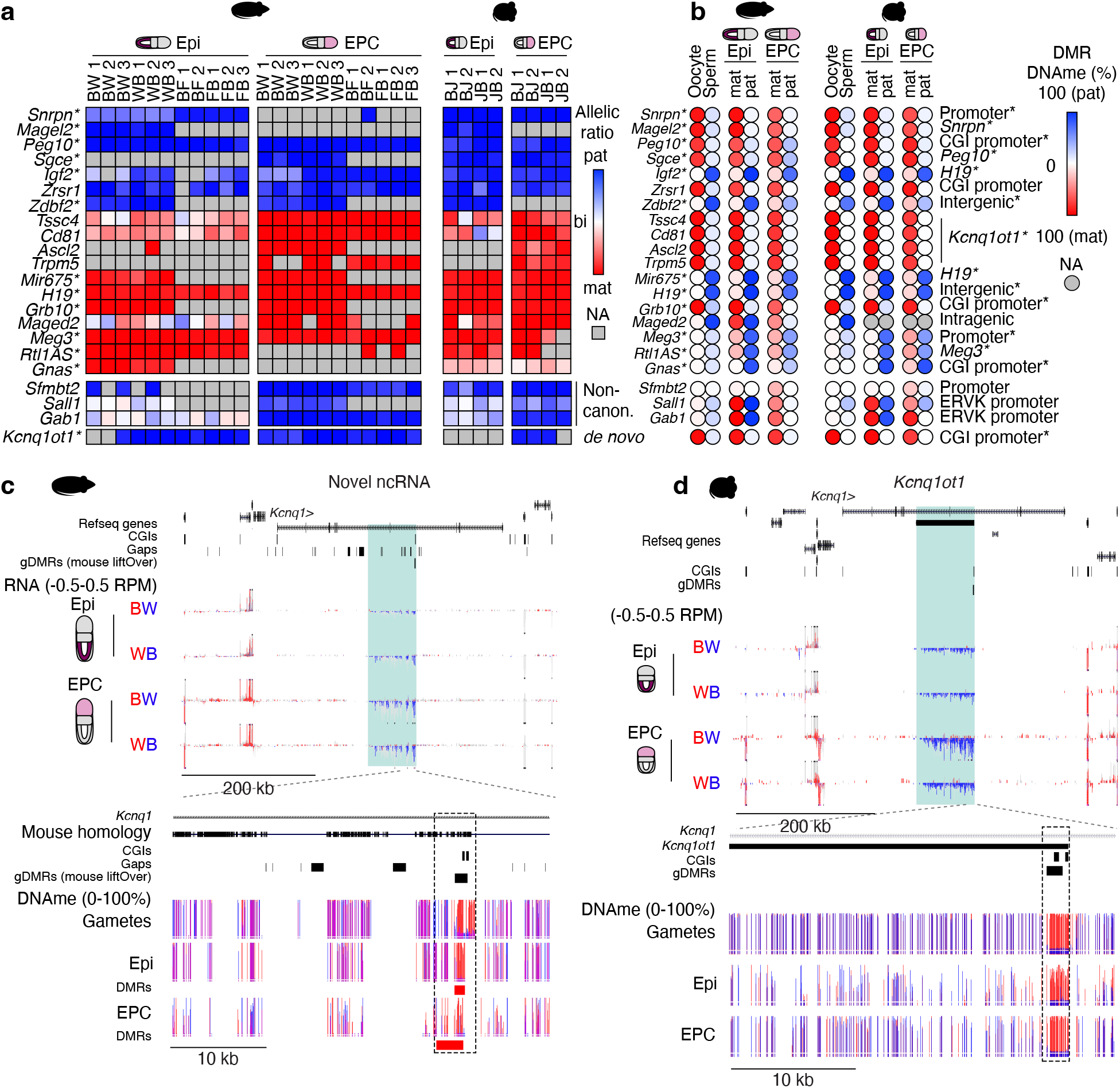
Conservation of genomic imprinting in rat and mouse. **a** Heatmap of parental expression ratios in rat and mouse Epi and EPC cells. Genes previously identified as imprinted in human are indicated with an asterisk. **b** Heatmap of allele-specific DNAme levels over DMRs associated with imprinted genes in **a**. The relative position of DMRs is included. DMRs that also show parent-specific methylation in human are indicated with an asterisk. **c-e** Rat and mouse genome browser screenshots of the *Kcnq1* locus. Figure legend as in **Fig. 1c**. In rat, a paternally expressed unannotated antisense ncRNA is expressed from the gene body of *Kcnq1*. The inlet tracks are ordered as follows: mouse homology, CGIs, parental rat strain SNVs and INDELs, gaps in the rat reference genome (rn6) and coordinates of mouse gametic DMRs projected onto the rat reference. DNAme levels are shown as bar plots, and CpGs covered by at least 1 allele-specific read (Epi, EPC) or 5 reads (gametes) are shown. Pink indicates methylation of both alleles. DMRs identified in rat are included. The relative position of the *Kcnq1ot1* CGI promoter DMR is indicated by a dashed box.

Among the canonically imprinted genes identified in rat, the growth-factor receptor binding gene *Grb10* shows maternal-biased expression in the epiblast and EPC and is associated with a maternally methylated DMR established in the oocyte (**Fig. 2a-b & Sup. fig. 3a**). The evolutionary conservation of maternal expression of *Grb10* is of particular interest as this gene shows maternal expression in mouse (48), confirmed here (*Grb10*:CGI promoter), and isoform-and tissue-specific imprinting in human (49). While there is only one annotated isoform of *Grb10* in rat, we find that conservation of oocyte-specific DNAme at the CGI promoter is maintained in embryonic and extra-embryonic tissues associated with maternal-specific expression in both cell types. *De novo* transcriptome assembly, described in detail below, clearly revealed two major *Grb10* isoforms in the rat, both of which are transcribed from the maternal allele in epiblast and EPCs (**Sup. fig. 3a**), consistent with what is observed in mouse.

Another canonically imprinted gene, the CCCH zinc finger gene *Zrsr1*, is paternally expressed in association with a gametic maternally methylated DMR in mouse (50), confirmed here (*Zrsr1*:CGI promoter) (**Fig. 2a-b, Sup. fig. 3b**). The *Zrsr1* gene is intronic and antisense to *Commd1*, a maternally expressed gene, and the maternally methylated *Zrsr1*:CGI promoter DMR is established in oocytes via *Commd1* transcription in mouse (51). Of note, mouse, rat and human all express *Commd1* at high (RPKM>15) levels in oocytes (**Sup. fig. 3c**). However, *Commd1* is biallelically expressed in human and is distal to *Zrsr1* (**Sup. fig. 3d**). Here, we show *Zrsr1* is paternally expressed in association with a maternally methylated *Zrsr1*:CGI promoter DMR in both mouse and rat (**Fig. 2a-b**). Thus, transcription of *Commd1* in oocytes, coupled with the proximal positioning in murids of *Zrsr1* and its CGI promoter, likely potentiated the co-evolution of *Zrsr1* and *Commd1* imprinting.

Surprisingly, another canonically imprinted transcript in mouse and human, *Kcnq1ot1*, was not identified in our allele-specific analysis. *Kcnq1ot1* is a paternally-expressed non-coding gene that is associated with silencing of adjacent genes, including *Kcnq1, Ascl2* and *Cd81* (45– 47), resulting in their maternal-specific expression. Despite the apparent lack of *Kcnq1ot1* in the rat, we confirmed maternal-specific expression of four orthologous imprinted genes: *Ascl2, Cd81, Trpm5* and *Tssc4* (**Fig. 2a**), prompting us to explore this genomic region in greater detail. Manual inspection revealed that *Kcnq1ot1* is not annotated in the rat reference genome “rn6” and thus was not assayed by our pipeline. Nevertheless, there was a clear paternal-specific RNAseq signal in the gene body of *Kcnq1*, as in mouse. To define *Kcnq1ot1* in the rat and other unannotated imprinted genes, we performed *de novo* transcriptome assembly of rat epiblast and EPC RNAseq data, agnostic to allelic assignment. This analysis uncovered an anti-sense gene within the gene body of rat *Kcnq1* (**Fig. 2c**). Importantly, this transcript, which shares 87% homology with mouse, is transcribed exclusively from the paternal allele in rat epiblast and EPCs (**Fig. 2a,c**). Furthermore, mirroring the mouse *Kcnq1ot1*:CGI promoter DMR (**Fig. 2d**), the CGI promoter of the putative rat *Kcnq1ot1* transcript is hypomethylated in sperm and hypermethylated in oocytes, and maintenance of parental DNAme levels is observed in epiblast and EPCs (**Fig 2b-c**). Indeed, agnostic identification of all DMRs between parental alleles identified the rat *Kcnq1ot1* CGI promoter in both epiblast and EPCs (**Fig. 2c**). The recently released “rn7” rat reference genome includes a novel 32-kb ncRNA identified by the automated NCBI Eukaryotic Genome Annotation Pipeline as *LOC120099961* and manual inspection confirms this novel transcript is likely the orthologue of mouse *Kcnq1ot1*. Thus, in addition to *in silico* prediction approaches, allele-specific RNAseq and WGBS can be leveraged to identify imprinted ncRNAs that are not annotated in the reference genome.

We next wished to determine whether non-canonical imprinted gene expression is evolutionarily conserved (22). Focusing on EPC-specific imprints, we found that three genes previously identified as imprinted in mouse: *Sfmbt2, Gab1* and *Sall1* (12, 15, 18), show both paternal expression (**Fig. 2a**) as well as an associated maternally methylated DMR (**Fig. 2b**) in rat and mouse EPCs. A fourth imprinted gene that is regulated by both maternal H3K27me3 and DNAme in mouse, *Slc38a4* (18), shows 81% paternal allele expression in rat but did not pass our stringent statistical cutoffs due to variability between replicates (**Sup. table 1**). Notably, the bobby sox homolog gene *Bbx*, one of the transiently imprinted genes associated with maternal H3K27me3 in mice (12, 13, 52), is clearly biallelically expressed in rat (**Sup. table 1**). Other non-canonical imprinted genes in mouse could not be assessed for allele-specific expression in rat due to a lack of parental genetic variation (*Jade1* and *Smoc1*) or the absence of an annotated ortholog (*Platr4* and *Platr20*). Interestingly, all three non-canonical imprinted genes identified in rat are not imprinted in human or macaques (53). Together, these data indicate that the establishment of non-canonical imprints at *Sfmbt2, Gab1* and *Sall1* likely originated in the rodent lineage. Furthermore, the maintenance of their imprinting status in both mouse and rat indicates that their dosage likely plays an important role in extraembryonic development, at least in murids.

### *De novo* identification of genomic imprinting in rat

While the application of cross-species comparisons of allele-specific expression and DNAme data led to the identification of 21 orthologous genes imprinted in rat and mouse, two factors constrain the comprehensive identification of imprinted genes using this approach. Firstly, the Ensembl gene annotation, which we relied on to analyze orthologous genes, includes many genes in the mouse with no apparent ortholog in the rat, including of known imprinted genes, such as *Kcnq1ot1* (**Fig. 2c**). If such orthologs are actually present in the rat genome, our reliance on Ensembl annotations precludes the comprehensive characterization of imprinting in the rat. Secondly, due to natural divergence, the rat may truly lack genes that are orthologs to those present in the mouse (such as *Platr4*), and vice versa. If so, novel imprinted genes in the latter category would not be identified using the approach described above.

To circumvent the shortcomings associated with identifying novel imprinted genes using homologous gene annotations (n=16,770 genes), we chose to use NCBI RefSeq transcript (n=69,157 transcripts) annotations to calculate total and parental genome-specific transcript expression levels, as reported above (**Fig. 1**). Additionally, since imprinted gene expression may arise from unannotated promoters (15), we supplemented the NCBI Refseq gene annotations with *de novo* transcriptome assembly using our epiblast and EPC RNAseq samples (n=3,242 additional transcripts). Using these new rat transcript annotations and employing the same statistical tests and filtering criteria described above, we identified 8 novel putative imprinted genes in the rat that do not show imprinting in the mouse in our data or in mouse imprinted gene databases (**Fig. 3a & Sup. fig. 4a-b**). An additional set of 33 genes that were scored as maternally expressed in rat EPCs were also found to be highly expressed (RPKM>1 from either allele) in adult blood (**Sup. fig. 4c-d**). Thus, all putative rat-specific imprinted genes identified here are imprinted in EPCs, in line with the observation in mouse and human that relative to embryonic or somatic tissues, the placenta exhibits a greater level of parent-of-origin specific gene expression. Furthermore, as none of these 8 genes are imprinted in human, their imprinting likely arose after the rat and mouse lineages diverged.

**Figure 3.**
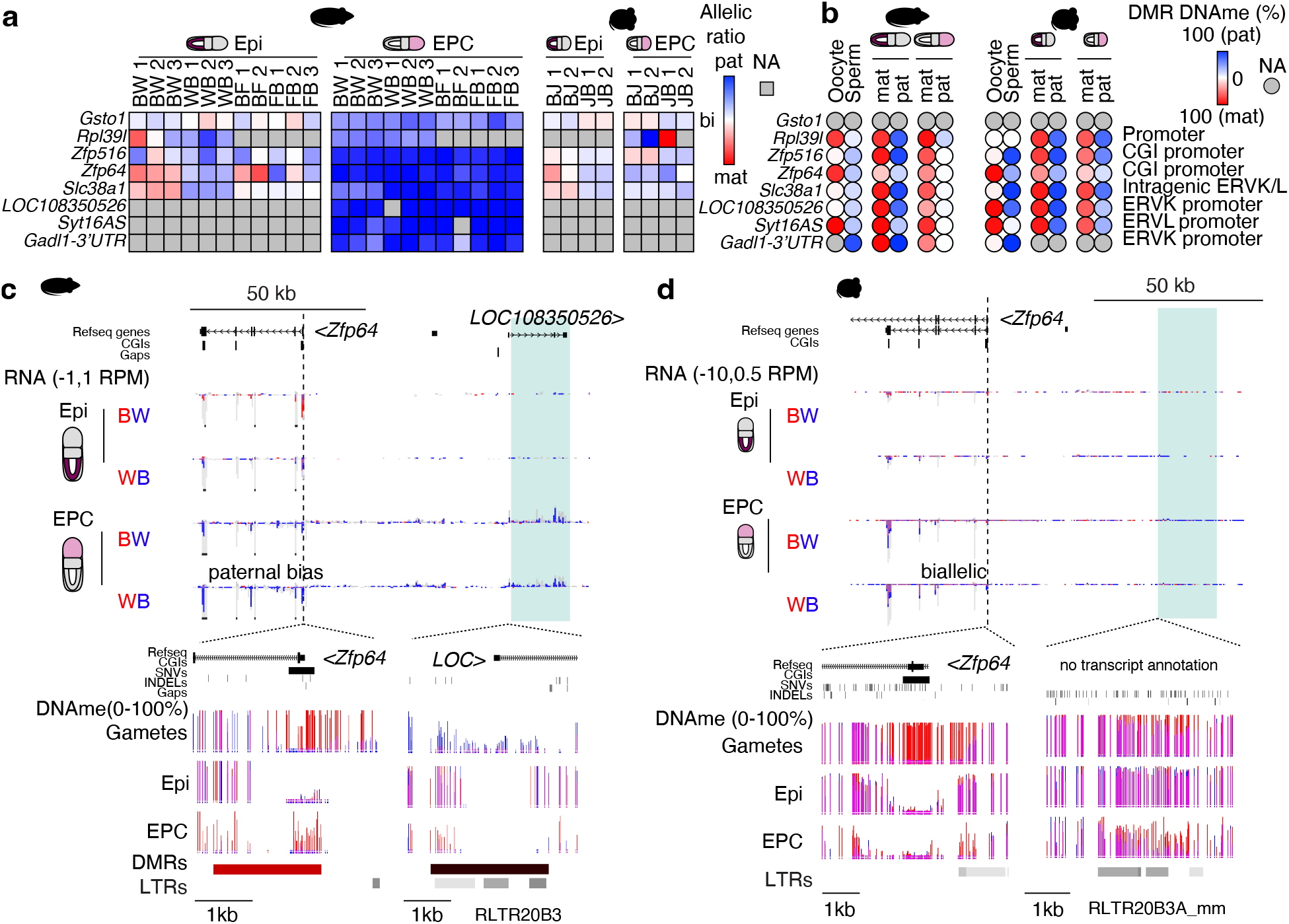
De novo genomic imprinting in rat. **a-b** Heatmaps of parental expression ratios and allele-specific DNAme levels in rat and mouse Epi and EPC cells as in **Fig. 2a-b**. Imprinted genes in rat that are not imprinted in mouse are shown. **c-d** Rat and mouse genome browser screenshots of the *Zfp64* and *LOC108350526* loci, two rat-specific imprinted genes. Browser tracks are as shown in **Fig. 2d**. The rat-specific gene *LOC108350526* and syntenic mouse region is highlighted in green. The location of LTR retrotransposons are included.

Consistent with non-canonical imprinting, post-fertilization DMRs are evident at 7/8 paternally expressed genes in rat EPCs, including *LOC108350526, Zfp64, Zfp516, Slc38a1, Gadl1-3’UTR, Syt16AS*, and *Rpl39l*. Another chromatin mark, such as maternal H3K27me3, may therefore be functioning as the imprint at these genes. As an example, the promoter of the novel rat gene *LOC108350526* shows rat-specific hypomethylation in oocytes and sperm (**Fig. 3b-c**). Consistent with non-canonical imprinting, DNAme on the maternal allele is acquired post-fertilization specifically in rat EPCs, resulting in asymmetric parental DNAme levels in association with paternal-biased expression (**Fig. 3b-c**). In mouse, the syntenic region of the rat *LOC108350526* promoter is hypermethylated and remains hypermethylated on both alleles following fertilization (**Fig. 3b,d**). This transcript has no homologue in mouse or human, and its putative open reading frame does not encode any known protein domain, suggesting it is a novel non-canonical imprinted ncRNA that arose in the rat lineage. Interestingly, the nearby gene *Zfp64* shows clear paternal expression exclusively in rat EPCs (**Fig. 3a,c**). The CGI promoter of *Zfp64* is maternally methylated in gametes in both rat and mouse, yet this parental asymmetry is maintained only in rat EPCs, as the locus becomes hypomethylated in the rat epiblast and mouse embryo (**Fig. 3b-d**). Consistent with the loss of DNAme, TET1, a methyl-cytosine dioxygenase (54), is enriched at the *Zfp64* CGI promoter in mouse ESCs (**Sup. fig. 4e**), perhaps explaining the loss of DNAme at this region in the mouse. The fact that *LOC108350526* is located adjacent to *Zfp64* makes this locus worthy of future investigations into the relationship between canonical and non-canonical imprinting (see Discussion). Taken together, these results indicate that canonical genomic imprinting generally shows a greater degree of conservation in mammals than does non-canonical imprinting, raising the question, what drives the establishment of species-specific non-canonical imprinting?

### Epigenetic profiling of rat oocytes

A common feature of the rat-specific imprinted genes identified here is paternal-specific transcription in extraembryonic tissue in the absence of a canonical DNA methylation imprint in oocytes. These are hallmark features of non-canonical imprinted genes in the mouse, which were recently shown to depend upon H3K27me3 deposited by PRC2 in oocytes (12, 13). As noted previously, rat oocytes show relatively fewer hypermethylated loci than mouse oocytes (**Sup. fig. 5a**) (39), perhaps enabling more widespread H3K27me3 deposition. We therefore hypothesized that rat-specific non-canonical imprinted genes are associated with rat-specific H3K27me3 domains in oocytes. To determine whether H3K27me3 is indeed enriched at rat-specific non-canonical imprinted loci, we profiled fully grown rat oocytes (FGOs) by CUT&RUN (55). Additionally, we profiled H3K4me3 and H3K36me3, which generally mark the promoter regions and gene bodies, respectively, of actively transcribed genes. Genome-wide analysis revealed strong correlation between biological replicates, as well as between H3K36me3 and DNAme levels, as shown previously in mouse oocytes (39) (**Sup. fig. 5b**). Additionally, we observed an anticorrelation between H3K27me3 and H3K36me3 in rat oocytes (**Fig. 4a**), consistent with the observation that H3K36me3 inhibits PRC2 activity *in vitro* (56, 57) and is anti-correlated with H3K27me3 *in vivo* (58). Chromatin state analysis using ChromHMM (59) revealed that H3K27me3 indeed marks a larger fraction of the genome in rat than mouse oocytes (**Sup. fig. 5c**). Conversely, H3K36me3 domains are more widespread in the mouse, likely reflecting a greater prevalence of non-genic transcripts in the mouse (39). Analysis of recently published H3K27me3 CUT&RUN data from rat oocytes (60) reveals a strong correlation with our data (Spearman rank correlation 0.95, **Sup. fig. 5d**), confirming the reproducibility of this method. Additionally, rat oocyte and early embryo H3K27me3 levels are generally correlated (Spearman rank correlation >0.67, **Sup. fig. 5d**), indicating that H3K27me3 levels are largely maintained in the rat embryo.

**Figure 4.**
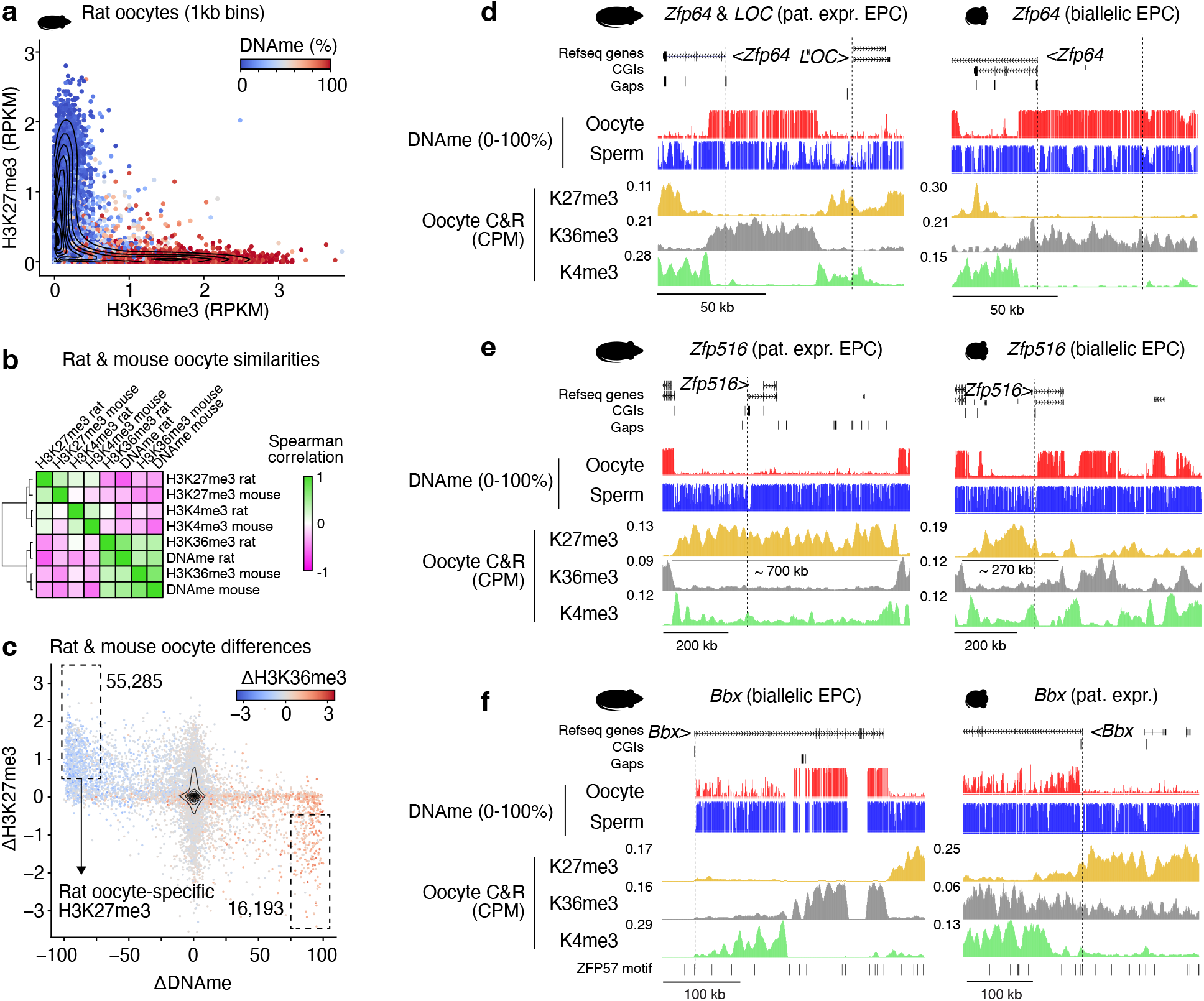
Species-specific H3K27me3 in oocytes is associated with species-specific non-canonical imprinting *Zfp64* and *Zfp516*, and transient imprinting of *Bbx*. **a** 2D scatterplot showing genome-wide H3K36me3 and H3K27me3 levels over 1kb bins in rat oocytes. Datapoints are coloured by average DNAme levels. A random subset of 10,000 bins is shown. The density of all data points (n=1,164,268) is summarized by a contour plot. **b** Cross-species Spearman correlation metrics between H3K4me3, H3K27me3 and H3K36me3, and DNAme levels over syntenic 1kb bins in rat and mouse oocytes (n=823,926). **c** Scatterplot showing differences in rat and mouse oocyte DNAme, H3K27me3 and H3K36me3 levels over syntenic 1kb bins. A random set of 10,000 bins are shown, and all bins (n=823,926) are summarized by a contour plot. The number of bins with a delta H3K27me3 >0.5 RPKM and DNAme >75% are indicated by a dashed box. **d-f** Rat and mouse genome browser screenshots of the *Zfp64*-*LOC108350526, Zfp516* and *Bbx* loci. Replicates of CUT&RUN data were merged and mean levels are shown as counts per million aligned reads (CPM). Promoters are indicated by a dashed line.

To directly compare rat and mouse oocyte epigenomes, we first assessed the enrichment of H3K27me3, H3K36me3, H3K4me3 and DNAme levels across syntenic 1kb regions, which revealed globally similar distributions of each of these marks (**Fig. 4b**). Indeed, genes expressed in rat and mouse oocytes show similar patterns of H3K4me3 and H3K36me3 over their promoters and gene bodies, respectively (**Sup. fig. 6a**). However, in mouse, the gene bodies of silent genes were also enriched for H3K4me3 (**Sup. fig. 6a**). Such “broad” H3K4me3 domains were previously identified in mouse oocytes (61–63) but were not observed in human oocytes (64), raising the question of the evolutionary origins and biological relevance of such domains. Rat and human H3K4me3 domains are on average 10kb (37%) smaller than those in mouse (**Sup. fig. 6b**), including at the previously analyzed *Kdm4c* promoter (**Sup. fig. 6c**). These results indicate that, with respect to H3K4me3, rat oocytes may be more similar to human than mouse. Consistent with the ChromHMM results and the relatively restricted distribution of H3K4me3, H3K27me3-enriched regions were >3x more abundant in rat than mouse (as measured in syntenic 1kb bins, **Fig. 4c**). Why the distribution of these chromatin marks differs so dramatically between these closely related species is an intriguing question that awaits further investigation.

We next focused on the epigenetic state of *Gab1, Sall1* and *Sfmbt2*, the three non-canonical imprinted genes showing paternal allele-specific expression in both the rat and mouse (**Fig. 2a**). Interestingly, all maternally methylated DMRs associated with these loci overlap with an H3K27me3 domain in oocytes of both murids (**Sup. fig. 7a**), confirming that non-canonical imprinting of these genes is conserved. In line with our findings, H3K27me3 is clearly maintained at these loci to the blastocyst stage in mouse and rat (**Sup. fig. 7a**). As recent reports have indicated that G9A and GLP play an important role in establishment (17) and maintenance (17, 18) of non-canonical imprinting, we analyzed our recent mouse oocyte H3K9me2 ChIP-seq data (65) and found that *Gab1, Sfmbt2* and *Sall1* also overlap an H3K9me2 domain (**Sup. fig. 7b**). Taken together, these data are consistent with the model that PRC2 and/or G9A/GLP play an important role in the establishment of non-canonical imprints.

Unlike *Gab1, Sall1* and *Sfmbt2*, the 8 genes mentioned above (**Fig. 3a-b**) are not imprinted in mouse, indicating that whatever the mechanism of imprinting establishment at these loci, it occurred relatively recently (<13 million years ago) in the rat lineage. Intriguingly, only *LOC108350526* showed relatively higher levels of H3K27me3 and concomitant lower levels of H3K36me3 and DNAme in rat compared to mouse oocytes (**Sup. fig. 8a**). Indeed, a rat-specific H3K27me3 domain that overlaps with the promoter of *LOC108350526* is clearly present in rat but absent over syntenic regions in mouse oocytes (**Fig. 4d**). Furthermore, while a maternal H3K27me3 domain exists at the *Zfp516* locus in both rat and mouse oocytes, the mouse H3K27me3 domain is adjacent to the *Zfp516* promoter and covers a region 2-3X smaller than in the rat (**Fig. 4e**). More specifically, the mouse H3K27me3 domain terminates at the gene body of *Zfp516*, perhaps as a consequence of transcription-coupled H3K36me3 deposition in the mouse oocyte. Indeed, *Zfp516* is expressed in mouse (RPKM = 3.07) but not rat (RPKM < 0.00) oocytes (**Sup. fig. 8b**). Thus, differential expression of *Zfp516* in rat and mouse oocytes correlates with a rat-specific H3K27me3 domain, which in turn may regulate imprinted expression in rat EPCs. Both *LOC108350526* and *Zfp516* maintain H3K27me3 levels to the blastocyst stage in rat, demonstrating that these are bona fide non-canonical imprinted genes (**Sup. fig. 8c**). Conversely, *Bbx* is a transiently imprinted gene associated with maternal H3K27me3 in mouse (12, 13, 52) that is biallelically expressed in rat (**Sup. table 1**). Comparison of the rat and mouse *Bbx* locus clearly reveals a mouse-specific H3K27me3 domain that overlaps with the CGI promoter in oocytes (**Fig. 4f**). Taken together, these observations indicate that the species-specific imprinting of a subset of genes in the mouse or rat may be explained by the species-specific targeting of PRC2 in oocytes. Analysis of the remaining rat specific non-canonical imprinted genes did not reveal a difference in H3K27me3 levels in oocytes, indicating that alternative epigenetic marks, and/or timing of H3K27me3 removal, is likely at play. For example, differences in H3K27me3 maintenance in early rat and mouse embryos could give rise to species-specific imprinting. However, comparing rat and mouse preimplantation embryo H3K27me3 data did not uncover any clear species-specific maintenance over the 8 rat-specific non-canonical imprinted genes identified here. Only one such gene, *Gsto1*, displayed partial loss of H3K27me3 levels at the 2C stage in mouse compared to robust maintenance to the blastocyst in rat (**Sup. fig. 8c**). Additional investigations into the dynamics of H3K27me3, H3K9me2 and other repressive marks in mammalian embryos, combined with functional studies, will hopefully provide insights into the relative importance of such repressive covalent histone marks in the establishment and maintenance of extraembryonic non-canonical imprinting.

Genetic differences can also contribute to species-specific imprinting. The *Slc38a1* gene, which encodes an amino acid transporter, is another putative rat-specific non-canonical imprinted gene (**Fig. 3a**). As an H3K27me3 domain overlaps the *Slc38a1* promoter in both rat and mouse oocytes and early embryos (**Sup fig. 8c**), H3K27me3 per se is apparently not sufficient to confer non-canonical imprinting. Analysis of DNAme levels on the other hand reveals a rat-specific maternally methylated DMR (*Slc38a1*:intragenic ERVL/K) in EPCs (**Fig. 3b**) that overlaps with an MTD retroelement (**Fig. 5a-b**). Analysis of sequence synteny uncovered two key aspects that differentiate the rat *Slc38a1* locus from that in the mouse. First, while the underlying intronic DMR sequence is present in mouse, only the rat gene harbours a ZFP57 binding motif in the differentially methylated MTD retroelement (**Fig. 5b**). Analysis of the MTD consensus sequence (66) reveals that the rat MTD element gained two substitutions, resulting in the genesis of a ZFP57 motif (GC**A**GC**G** --> GCGGCA). Interestingly, 7/39 distinct rat strains have a substitution (GCG**A**CA) “away” from the MTD consensus that abrogates the ZFP57 motif (67). While the homologous sequence in mouse (GCAGCA) shares one substitution with the rat, this sequence does not conform to a ZFP57 motif. No further substitutions occurred over those 6 bases in any of the 36 sequenced mouse strains (68), including JF1 (69). Secondly, the canonical CGI promoter of *Slc38a1* contains a rat-specific ERVK promoter insertion (**Fig. 5b**). Whether either of these features are causal for the establishment of non-canonical imprinting remains to be tested, perhaps by analyzing rat strains that subsequently lost the ZFP57 motif. As maintenance of differential gametic DNAme at loci showing canonical imprinting is dependent upon ZFP57/ZFP445 binding, it is tempting to speculate that the novel ZFP57 motif in the intron of *Slc38a1* plays a role in its non-canonical imprinting. Since rat gametes are unmethylated at the novel ZFP57 motif, an allele-specific gain of maternal DNAme must occur post-fertilization, perhaps in conjuction with loss of maternal H3K27me3 (14). Further studies aimed at understanding how maternal H3K27me3 established in oocytes is replaced by maternal-specific methylation in extra-embryonic tissues will deepen our understanding of the relationship between these repressive epigenetic marks.

**Figure 5.**
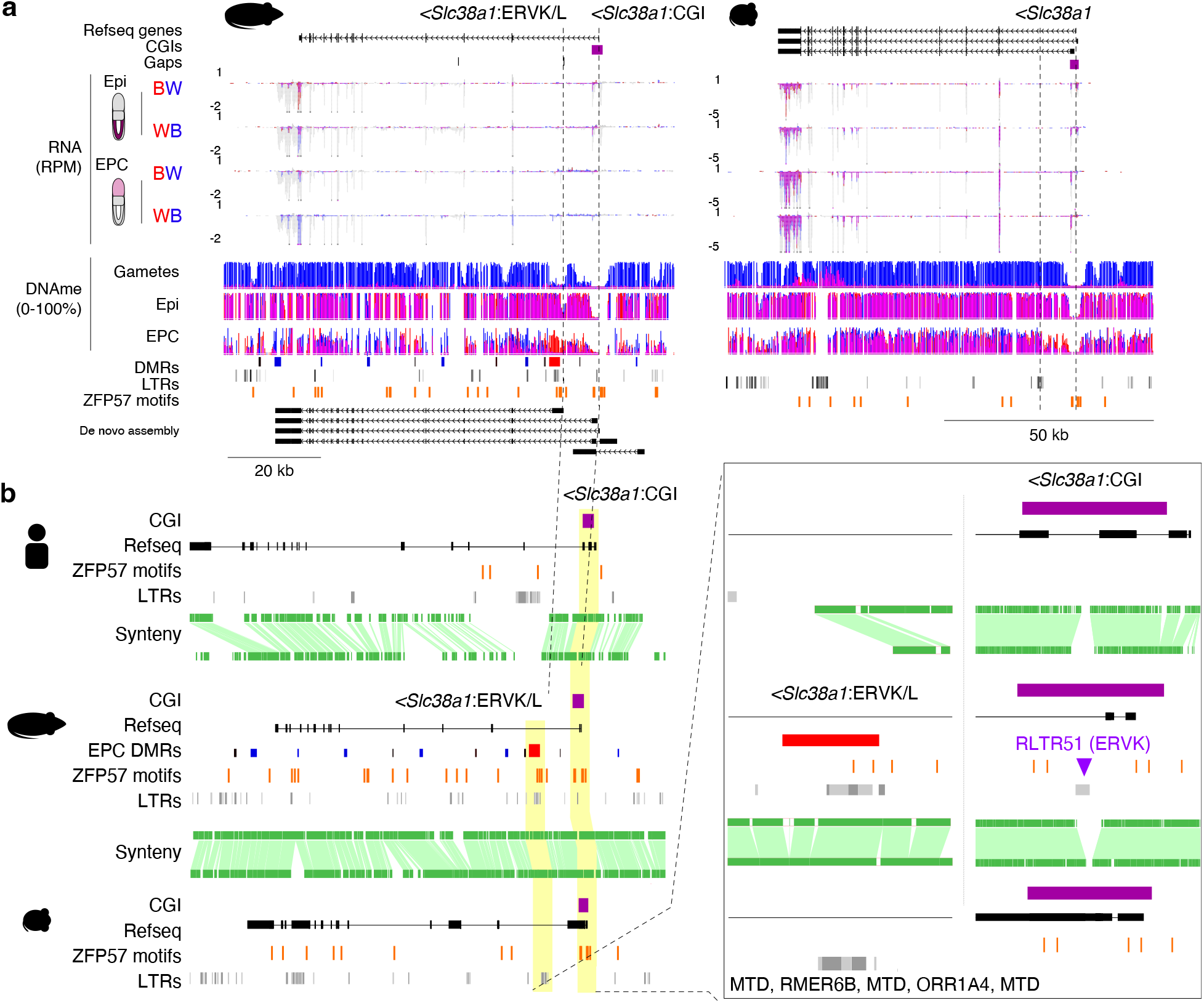
Rat-specific non-canonical imprinted gene *Slc38a1*. **a** Rat and mouse genome browser screenshots of the *Slc38a1* locus. *Slc38a1* is paternally expressed in rat and biallelically expressed in mouse EPCs. Tracks are presented as in **Fig. 2d b** Ensembl Region Comparison screenshot of the *Slc38a1* locus in rat (rn6), mouse (mm10) and human (hg19). Syntenic regions are shown in green. The locations Refseq genes, LTRs, CGIs (purple) and ZFP57 binding motifs (orange) are shown. Rat EPC DMRs are included. The CGI promoter and intronic DMR are highlighted in yellow and magnified (right panel) for clarity.

### Alternate promoters of murid-specific non-canonical imprinted genes

Interestingly, maternally methylated DMRs that arise following fertilization frequently overlap alternative promoters derived from ancient insertions of endogenous retroviruses (15), which harbour strong promoters in their long terminal repeats (LTRs). To assess the potential evolutionary conservation of LTRs in promoting imprinted gene expression, we analyzed syntenic regions between rat, mouse and human. For example, the non-canonical imprinted gene *Gab1* is expressed from the paternal allele in rat and mouse EPCs, in association with an EPC-specific DMR established post-fertilization (**Sup. fig. 9a**). Importantly, the *Gab1* DMR overlaps an alternate ERVK LTR promoter that drives paternal-specific expression of *Gab1* in both species (**Sup. fig. 9b**). Notably, this ERVK LTR is absent from the human genome, implicating the insertion of this ERVK element in the rodent lineage in the provenance of imprinted expression of *Gab1*. Similar to *Gab1*, a maternal H3K27me3 domain also overlaps the *Sall1* promoter in rat and mouse oocytes (**Sup. fig. 9c**), and an upstream ERVK promoter present in rat and mouse but absent in the orthologous human locus overlaps an extra-embryonic DMR and drives paternal-specific expression of *Sall1* in EPCs (**Sup. fig. 9d**). Interestingly, *Sall1* was predicted to be imprinted in human using in silico approaches (70), yet no empirical data has yet been produced to support this prediction. In contrast, while *Sfmbt2* shows imprinted gene expression in mouse and rat EPCs and overlaps a maternally deposited H3K27me3 domain (**Sup. fig. 9e**), the promoter of this gene does not overlap an annotated LTR element (**Sup. fig. 9f**). As five gaps exist in the rat reference genome between the *Sfmbt2* CGI promoter and the putative RLTR11B ERVK alternate promoter identified in mouse (15), we cannot rule out the possibility that an alternate promoter embedded in this region contributes to the imprinted expression of *Sfmbt2* in the rat. Taken together, these results are consistent with the model that alternate promoters, particularly those provided by LTRs, play an important role in promoting non-canonical imprinted expression of genes showing murid-specific imprints, and that maternally deposited H3K27me3 likely serves as the gametically inherited genomic imprint at many of these loci.

### Imprinted X chromosome inactivation in rat

X chromosome inactivation (XCI) involves the random and widespread silencing of genes on one X chromosome in adult female cells. During mouse embryogenesis however, XCI is not always random, with the fidelity of XCI influenced by genetic background. Indeed strain-specific XCI is observed across most adult mouse tissues (71). Furthermore, placental cells preferentially inactivate the paternally inherited X chromosome in mice (25, 72), likely due to non-canonical imprinting of *Xist* (12, 13, 73, 73). Interestingly, a recent study showed that H3K27me3 is enriched at the *Xist* gene in rat oocytes and early embryos (60), consistent with cytological evidence of non-random XCI in yolk sacs (74) and our CUT&RUN data (**Fig. 6a**). To determine whether either mode of XCI skewing occur in rat, we calculated the mean paternal expression ratio of all autosomal and X-linked transcripts (**Fig. 6b**). Autosomal transcripts are mostly biallelically expressed in epiblasts and EPCs, indicating that maternal decidua contamination is minimal. Rat epiblast cells do not show XCI skewing, suggesting that rat lab strains have not undergone sufficient sequence divergence to favour allelic bias in XCI. However, in line with findings in mouse, the paternally inherited X chromosome is preferentially inactivated in EPCs. Thus, imprinted XCI in the trophoblast lineage is conserved in rat and mouse.

**Figure 6.**
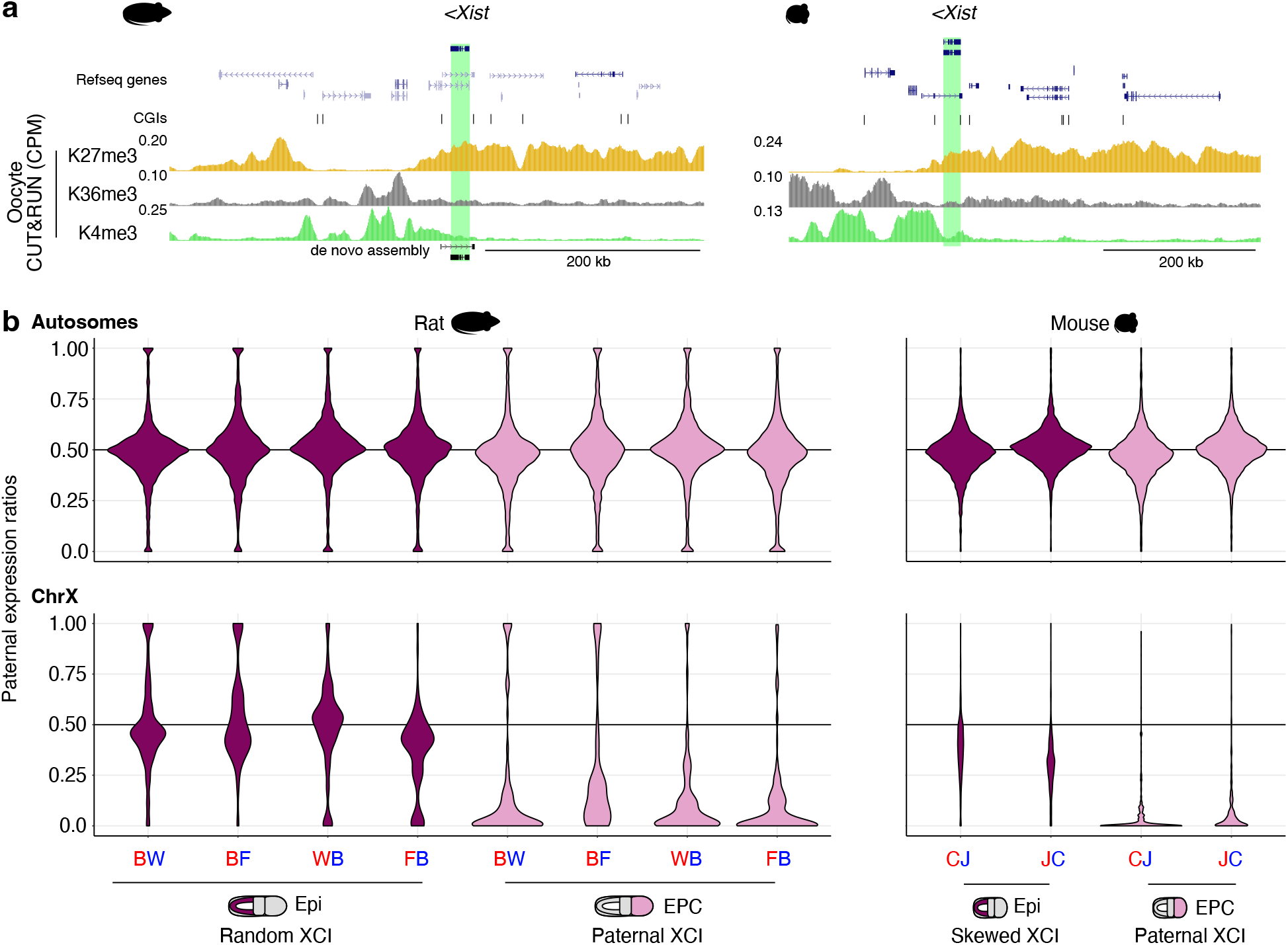
X chromosome inactivation (XCI) in rat and mouse. **a** Rat and mouse genome browser screenshots of the *Xist* locus. Refseq genes and CGIs are included. CUT&RUN (rat) and ChIP-seq (mouse) data are shown in counts per million (CPM). *Xist* is highlighted in green. **b** Violin plots showing the distribution of paternal expression ratios of autosomal (top) and X-chromosome (bottom) in rat and mouse epiblast and EPC samples. Replicate 1 of mouse cross “CJ” was omitted due to an XO genotype.

## DISCUSSION

The rat is an increasingly important model organism, yet few studies have focused on genomic imprinting in this species despite the importance of imprinted gene regulation for proper development, behaviour and physiology. Indeed, only 13 imprinted genes have been described thus far in rat, all of which were previously identified in mouse or human (11, 34, 60, 75). Here, employing a genome-wide approach, we expand this list to 19 canonical and 11 non-canonical imprinted genes. We further characterize the epigenetic marks present at these loci in oocytes as well as early embryonic tissues, and their association with parent-of-origin regulated gene expression, thereby providing a comprehensive map of the rat imprintome and the potential mechanisms by which imprinting is established.

Comparative analyses of imprinting in mice and humans indicate that the majority of canonically imprinted genes are conserved between rodents and primates, which diverged ∼90 million years ago (76). In contrast, we report the non-canonical imprinting of several genes in Muridae, including *Sfmbt2, Gab1* and *Sall1*, that were not identified in recent screens of imprinting in humans (77) or macaques (53). While we identified H3K27me3 deposited in oocytes as a putative imprint of these three genes, further studies are required to determine whether H3K27me3 is essential for the establishment of their imprinting in rat. In addition to PRC2, G9A and GLP were recently implicated in the establishment (17) and maintenance (17, 18) of non-canonical imprinting in mouse. Indeed, imprinted expression of *Gab1, Sfmbt2* and *Sall1* was shown to depend on both H3K9me2 and H3K27me3 pathways and all three genes overlap an H3K9me2 domain in oocytes (65). While G9A has low H3K27 methyltransferase activity *in vitro* (78), and ancestral SET-domain proteins in Paramecium show both H3K9 and H3K27 methyltransferase activity *in vivo* (79), experiments on the molecular basis of non-canonical imprinting revealed that imprinted expression is disrupted in *Eed* maternal knock-out embryos, coincident with loss of maternal H3K27me3 (13). Taken together, these observations indicate that H3K9 and H3K27 methylation are likely deposited by independent pathways and function in a non-redundant manner. Whether establishment and maintenance of the 8 non-canonically imprinted genes unique to the rat are subject to the same imprinting pathways remains to be determined. Interestingly, recent studies profiling H3K27me3 levels in human, macaque, cow, pig, rat and mouse embryos demonstrated that this mark is maintained following fertilization only in rat and mouse (53, 60). Whether an alternate repressive pathway, such as G9A and GLP, compensates for the maintained repression of the maternal allele of non-canonically imprinted genes in human, cow and pig remains to be determined. Furthermore, whether the imprinted expression of rat-specific imprinted genes contribute to differences in rat and mouse physiology and/or behaviour remains to be established.

Species-specific imprinted gene expression specifically in extra-embryonic tissue supports the model that the developing placenta is at the nexus of ongoing “parental-conflict” in gene dosage (23). The most compelling evidence for the parental conflict hypothesis is based on functional analyses of imprinted genes in the mouse, which reveal that many such genes play a role in regulating embryo and offspring size (80). Among the novel imprinted genes identified in the rat, *Zfp516* (79% and 51% expressed from the paternal allele in EPC and epiblast cells, respectively) is of particular interest. The mouse orthologue is biallelically expressed (51% paternal allele expression) in epiblast and EPCs (**Sup. table 1**) and is essential for embryogenesis and brown adipose tissue (BAT) production (81). *Zfp516* binds and induces transcription of BAT genes, including Uncoupling Protein 1 (*Ucp1*), which in turn regulates a host of BAT-enriched genes (81) and is used as a biomarker for BAT in mouse and human (82). This is consistent with other imprinted genes linked to adipogenesis and fat metabolism, including *Mest, Klf14, Dlk1, H19, Peg3, Peg10, Grb10* and *Cdkn1c* (83). Thus, the recent evolution of *Zfp516* imprinted regulation in the rat lineage, wherein expression from the paternal allele promotes transcription and in turn brown fat production, including in the developing placenta, is consistent with the parental conflict theory.

The rat-specific imprinting status of *Slc38a1* is also noteworthy as four other genes within the same gene family, *Slc38a4, Slc22a18, Slc22a2* and *Slc22a3*, are imprinted in the mouse. All these genes encode related solute carriers, and *Slc38a4* is highly expressed in the placenta, where nutrient transport is critical. Indeed, *Slc38a4* knock-out mice have a placental phenotype affecting embryonic growth(84). Canonical imprinting of *Slc38a4* in mice is dependent upon transcription in oocytes of a murid-specific retroelement that inserted upstream of the *Slc38a4* CGI promoter (85), leading to *de novo* DNAme of the canonical DMR. Notably, *Slc38a4* is paternally expressed in the mouse placenta (15, 86) and G9A-depleted embryos fail to maintain the maternally methylated DMR (87), suggesting that both canonical and non-canonical imprinting mechanisms govern *Slc38a4* expression. Thus, in line with the observation that imprinted genes converge on shared pathways (10), *Slc38a1* represents a strong candidate rat-specific non-canonical imprinted gene. Given the observed sequence-specific differences at the *Slc38a1* CGI promoter and intronic DMR, investigation of the relative roles of retroelement transcription in EPCs and ZFP57 binding motifs in maintenance of post-fertilization gain of maternal DNAme will likely yield important insights into the evolution of non-canonical imprinting.

*Slc38a1* and *Zfp64*, another rat-specific non-canonically imprinted gene, were recently identified as paternally expressed in association with maternal H3K27me3 in mouse blastocysts (52). However, the asymmetry in parental expression was lost following implantation in mouse, which may explain why these genes were not identified in previous studies focusing on post-implantation tissues such as EPCs. Nevertheless, transient imprinting of mouse *Slc38a1* and *Zfp64* is in line with the observation that global levels of maternal H3K27me3 are maintained to the blastocyst stage in murids (60). In other words, *Slc38a1* and *Zfp64* are seemingly “primed” for imprinting in rat and mouse, yet the maintenance of these imprints is specific to the rat. Further analysis of such transient (to the blastocyst stage) and non-canonical imprinted genes will serve as important models for studying the provenance of imprinting maintenance. Indeed, our meta-analysis of the *Slc38a1* locus implicate a gain of a putative ZFP57 binding site and/or an insertion of an alternate ERVK/L promoter in the rat genome.

*Zfp64* is a zinc-finger protein that regulates mesenchymal differentiation via Notch signaling in mouse (89). Additionally, *ZFP64* directly promotes the expression of *MLL* in human leukemia (90). While *Zfp64* was found to be imprinted in the mouse placenta (91), a recent extensive search for imprinted genes revealed only modest (<70%) paternal-biased expression in an array of mouse tissues (88). Here, we confirm these findings, and show that *Zfp64* is indeed biallelically expressed in embryonic (50% paternal expression) and extra-embryonic (63% paternal expression) mouse cells. Consistent with this observation, its CGI promoter is hypomethylated in both cell types. In contrast, the rat *Zfp64*:CGI promoter is maternally methylated in association with imprinted expression (95% paternal) in ectoplacental cone cells. The genomic imprint of the *Zfp64*:CGI promoter is conserved between rat and mouse, in association with active transcription of a murid-specific MTC LTR in oocytes (39). Notably, *Zfp64* has not been reported as imprinted in human. However, the *Zfp64* CGI promoter lacks ZFP57 binding motifs in both rat and mouse, and ZFP57 in mouse ESCs is not enriched at this region (data not shown), as determined by ChIPseq (92). Thus, rat-specific imprinting of *Zfp64* cannot be explained simply by these sequence differences. However, species-specific demethylation of the *Zfp64* CGI promoter could be due to differential activity of the TET enzymes at this locus. Consistent with this possibility, TET1 is enriched at the *Zfp64* CGI promoter in mouse ESCs. TET1 profiling in mouse and rat embryos would confirm whether differential active demethylation can result in species-specific imprinting, and yield insights into ZFP57-independent modes of imprint maintenance.

In contrast to the genetic differences between rat and mouse *Zfp64, Slc38a1* and *Slc22a3*, rat-specific imprinting of *Zfp516* and *LOC108350526* is associated with domains of H3K27me3 in oocytes unique to the rat. While the molecular basis of PRC2 targeting in oocytes remains to be determined, H2AK119 ubiquitination is apparently not required for maintenance of H3K27me3 or imprinted expression of non-canonical imprinted genes after fertilization (93, 94). While the mechanism underlying the switch from maternally inherited levels of H3K27me3 to maternal-specific dense DNA methylation in the extraembryonic lineage remains unknown (14, 15, 22), the paternally expressed gene Zinc Finger DBF-Type Containing 2 (*Zdbf2*) may offer insights. This gene is regulated by a maternally methylated somatic DMR that overlaps with a domain of H3K27me3 in oocytes and a H3K27me3-to-DNAme switch was shown to be induced by active transcription across the locus which, in turn, is promoted by an upstream paternally methylated gametic DMR (95). Indeed, genetic ablation of both DNMT1 and EZH2 results in biallelic *Zdbf2* expression in mouse zygotes (18). Whether a similar phenomenon is at play at all non-canonical imprints that undergo H3K27me3-to-DNAme switching remains to be tested.

## CONCLUSIONS

In summary, our results provide the first atlas of genomic imprinting in the rat, including 22 conserved and 8 novel candidate rat-specific imprinted genes. Notably, several of the novel genes identified are imprinted specifically in placental precursor cells and have been previously shown in other species to play an important role in metabolism. Determining whether disruption of such non-canonical imprinting impacts fetal and/or placental growth is an obvious avenue for future investigation. Clearly, measuring the extent and conservation of imprinted gene expression in additional mammals, such as pig, cow, human (60) and macaques (53), complemented by interrogation of genetic and epigenetic differences at species-specific imprinted loci, will deepen our understanding of the evolution of genomic imprinting. Finally, there is an increasing body of evidence implicating canonical imprinted genes in complex behavioural traits (10, 96–100). Our observation that canonical imprinted gene expression is highly conserved in the rat opens the door to future mechanistic studies on the role of these genes in behavioural and other traits in this increasingly tractable model.

## METHODS

### Rat and mouse embryo collection

E8.5 F1 hybrid rat embryos were isolated from reciprocally crossing BN/CrlCrlj (BN rat, Charles River) and WKY/NCrlCrlj (WKY rat, Charles River) strains as well as between strains BN/CrlCrlj and F344/NSlc (F344/N rat, Japan SLC). E7.25 F1 hybrid mouse embryos were isolated from reciprocally crossing C57BL/6N (Clea Japan) and JF1/Ms (National Institute of Genetics) strains. For both rat and mouse embryos, the ectoplacental cone (EPC) was dissected from the rest of the embryo before removing the visceral endoderm (VE). Finally, the epiblast (EB) was dissected from the extraembryonic ectoderm (ExE). Each ExE lysate was used for rapid genetic sex determination by direct PCR amplification of the Y chromosome gene using specific primers for mouse *Zfy* (5’-CCTATTGCATGGACTGCAGCTTATG-3’ and 5’-GACTAGACATGTCTTAACATCTGTCC-3’) and rat *Sry* (5’-CATCGAAGGGTTAAAGTGCCA-3’ and 5’-ATAGTGTGTAGGTTGTTGTCC-3’). Only female epiblasts and EPCs were subjected to strand-specific RNA-sequencing and whole genome bisulphite sequencing (in the case of BN-F344/N hybrids, only RNAsequencing was performed).

### Rat oocyte collection

GV stage rat oocytes were collected from ovary of 3-4 wk old BrlHan:WIST (Clea Japan) rats 48-50 h after administration of PMSG. Briefly, cumulus-oocyte complexes were collected in PB1 with 200 µM IBMX (Sigma), and subsequently incubated in the presence of 5-10 µg/ml cytochalasin B for 5-10 min to loosen the connection between cumulus cells and oocytes. Then, the cumulus cells were mechanically removed by repeated pipetting using a glass capillary, and the zona pellucida were removed by brief incubation in acidic Tyrode’s solution.

### Strand-specific RNA library preparation and sequencing

One nanogram of embryonic total RNA was reverse transcribed using SMARTer Stranded Total RNASeq Kit v2 -Pico Input Mammalian (Takara Bio) according to the manufacturer’s protocols, in which Read 2 corresponds to the sense strand by the template-switching reactions. The RNAseq libraries were quantified by qPCR using KAPA Library Quantification Kit (Kapa Biosystems). All libraries were mixed and subjected to paired-end 100 bp sequencing (paired-end 101 nt reads in which the first 1 nt of Read 1 and the last 1 nt of Read 2 were trimmed) on HiSeq 2500 system (Illumina). For each library, reads were trimmed using Trimmomatic (v0.32) (104) to remove 2 nucleotides of the 5’ end of read 2.

### Whole genome bisulphite sequencing (WGBS)

One hundred nanograms of genomic DNA from JF1 (female) and C57BL/6 (male) mouse hybrids were prepared for Methyl-seq (Illumina) library construction. Twenty nanograms of genomic DNA from the other mouse and rat hybrid crosses were prepared for rPBAT or tPBAT library construction. Each WGBS library was quantified using KAPA Library Quantification Kit (Kapa Biosystems). The tPBAT libraries from BN (female) and WKY (male) hybrids and Methyl-seq libraries were subjected to paired-end 150 bp sequencing on HiSeq-X-Ten system (Illumina). The other tPBAT and rPBAT libraries were subjected to paired-end 100 nt sequencing (paired-end 101 nt reads in which the first 1 nt of Read 1 and the last 1 nt of Read 2 were trimmed) on the HiSeq 2500 system.

### CUT and RUN library preparation and sequencing

CUT&RUN libraries were prepared as previously described (13) with some modifications. Briefly, oocytes were incubated with rabbit anti-H3K27me3 antibody (1/100, Diagenode, #C15410069), rabbit anti-H3K4me3 antibody (1/100, Active Motif, #39159), or rabbit anti-H3K36me3 (1/100, Abcam, #9050) in a wash buffer (20 mM HEPES (pH7.5), 150 mM NaCl, 0.5 mM Spermidine, complete EDTA-free protease inhibitor cocktail, 0.02% Digitonin) containing 2 mM EDTA at 4C on a plastic dish overnight. After the oocytes were washed twice by the wash buffer on a plastic dish, they were incubated with Protein A-MNase (500 ng/ml diluted in the wash buffer) provided by the Steven Henikoff Lab (55) at 4C for 3 hrs on a plastic dish. After washing twice, the oocytes were transferred into a 1.5 ml DNA LoBind tube (Eppendorf) containing pre-activated BioMag Plus Concanavalin A (ConA) beads (Bang Laboratories, Inc). The sequential step of the antibody incubation followed by the ConA binding prevents loss of oocytes during wash steps. After the oocytes were bound to ConA beads and the buffer was replaced to the wash buffer, CaCl2 was added at a final concentration of 2 mM to activate the MNase. After 20 min of digestion on ice, the digestion reaction was stopped, DNA fragments were recovered, and DNA libraries were prepared as described previously (13). PCR amplification was performed for 11-15 cycles. Libraries were sequenced on a NextSeq500 with single-end 75 bp reads (Illumina).

### RNAseq read alignment and allele-agnostic quantification using a haploid reference genome

Sequencing reads were trimmed using Trimmomatic (v0.32) and the following parameters: SLIDINGWINDOW:3:10 MINLEN:36 ILLUMINACLIP:TruSeq2-3-PE.fa:2:30:10. Read pairs that survived trimming were aligned to genome build mm10 or rn6 using STAR (v2.4.0.i) and PCR duplicate reads were flagged using Picard MarkDuplicates (v1.92). Library quality was assessed using samtools flagstat (v1.1) and Picard CollectRNASeqMetrics. Uniquely aligned, non-PCR-duplicate reads were kept for downstream analysis using samtools parameters: samtools view - bh -q 255 -F 1540. NCBI RefSeq mouse transcript annotations (n=106,520 transcripts, 35,977 genes) were downloaded from the UCSC Table Browser (last updated 2017-11-16). NCBI RefSeq rat transcript annotations (n=69,194 isoforms, 30,871 transcripts) were downloaded from the UCSC Table Browser (last updated 2018-03-09). Transcript expression values were calculated by averaging read coverage over exons using VisRseq (v0.9.12) and normalized to the total number of aligned reads and transcript length in kilobases (RPKM). Genome browser compatible normalized bigWigs were generated using bedtools genomecov (v2.22.1).

### Comparative analysis of mouse and rat homologous gene expression

Mouse and rat homologous gene annotations were downloaded from Ensembl Biomart (n=14,297). Mouse and rat epiblast and EPC gene expression similarity matrices and hierarchical clustering was subsequently generated using the Morpheus (Broad Institute) Spearman correlation tool on adjusted (log2(RPKM+1)) RPKM values. Mouse-rat syntenic 1kb bins (n= 1,899,360) were generated previously (39), and syntenic regions were illustrated using Emsembl Comparative Genomics and Region Comparison tools.

### Allele-specific RNAseq analysis using a diploid pseudogenome

Strain-specific SNVs and INDELs were downloaded from (3, 4, 68, 69), and diploid pseudogenomes “C57BL/6NJ x JF1” and “BN/Mcwi x WKY/NCrl” and “BN/Mcwi x F344/N” were generated using MEA. Allele-specific read alignment was performed using MEA with default parameters. Allele-specific transcripts expression over exons was calculated using VisRseq (v0.9.12) and the contribution of paternal-allele transcript expression was calculated using the formula:

Paternal bias = paternal coverage / (paternal + maternal coverage)

Strand-and allele-specific bigWigs were organized into UCSC Track Hubs along with reference-aligned bigWigs, as previously described, and used for screenshots over regions of interest.

### Identifying rat imprinted genes

NCBI RefSeq rat transcript annotations (n=69,194 transcripts, 30,871 genes) were downloaded from the UCSC Table Browser (last updated 2018-03-09), and total and allele-specific transcript expression was quantified as described above. An additional set of 3,242 transcripts were identified by *de novo* transcriptome assembly using Stringtie (v2.1.4) (105) and default parameters followed by merging with the NCBI Refseq transcript annotations using Stringtie -- merge and default parameters. Transcripts were filtered based on total (allele-agnostic) expression (RPKM ≥1) and allele-specific coverage (RPM ≥0.5). Paternal expression ratios were calculated as described above. Since transcripts are not necessarily expressed in all cell types analyzed in this study, or do not necessarily overlap a parental genetic variant, a paternal bias score of “-2” or “-1” was specified to transcripts not expressed (RPKM < 1) or did not meet allelic thresholds (allelic RPM <0.5), respectively. Student’s T-tests were performed on the paternal expression ratio value for each transcripts that passed filtering criteria in at least 1 sample, and Bonferroni correction was applied to the resulting P-values. Transcripts with an adjusted p-value <0.05 and for which sufficient allele-specific and total coverage was reported in at least 6 samples (rat) or 4 samples (mouse) were categorized as imprinted. It should be noted that since extra-embryonic tissues are located between maternal and embryonic tissues, extra-embryonic cell isolation can be contaminated with maternal decidua, which can in turn confound and overestimate the number and extent of maternally expressed imprinted genes (106). An additional filtering step was performed for rat EPC samples, whereby maternally expressed imprinted transcripts that are expressed in blood (RPKM >1 from PRJEB23955 (107), 9 replicates) were discarded. To avoid the artifactual scoring of maternally expressed transcripts caused by possible residual maternal decidua in rat EPC samples, we further categorized 35/42 maternally expressed imprinted transcripts as being normally expressed (allele-agnostic RPKM >1) in adult blood. Known imprinted genes in mouse such as *Ascl2, Trpm5* and *Tssc4* are included in the list of 35 genes, reflecting stringent annotations. Given their shared imprinted status in mouse, *Ascl2, Trpm5* and *Tssc4* are reported in **Fig. 1d** and **Fig. 2a-b**.

An additional test for imprinted expression in rat was carried out using linear modeling using Limma v3.50.1 (108) in R v4.1.2 (109). Transcripts with allelic RPM ≥0.5 on either allele in at least 2 replicates in both crosses (BWxWB and BFxFB) in either tissue (epiblast or EPC) type were kept. An eBayes statistic was calculated following linear model fitting using “lmFit”, and transcripts with an adjusted p-value <0.05 and a ≥4-fold change in expression between parental alleles were kept. The same filtering step for maternally expressed imprinted genes in EPC samples described above was applied. A set of 37 imprinted genes were identified, 16 of which were not identified in the T-test described above, and were included in **Fig. 1d**. Two genes identified by Limma, *Itga1* and *LOC103691708*, are encoded on chromosome 2 and are not included in **Fig. 1d** due to space limitations. See **Sup. Table 1** for the full list of imprinted genes identified by T-tests and Limma.

### Identifying mouse imprinted genes

NCBI RefSeq mouse transcript annotations (n=106,520 transcripts, 35,977 genes) were downloaded from the UCSC Table Browser (last updated 2017-11-16) and transcript expression was quantified as described above in rat.

### Agnostic identification of rat parent-specific DNA methylation

Sequencing reads were trimmed using Trimmomatic (v0.32) and the following parameters: SLIDINGWINDOW:3:10 MINLEN:36 ILLUMINACLIP:TruSeq2-3-PE.fa:2:30:10. Read pairs that survived trimming were aligned to diploid rat or mouse pseudogenomes (as described above) using MEA and Bismark (v0.16.3). Duplicate reads were removed and the number of methylated and unmethylated reads overlapping each CpG of the rat or mouse genome was reported. This resulted in total (allele-agnostic), maternal- and paternal-specific CpG report files. Replicates and reciprocal crosses were merged for visualization and analyses. CpGs covered by at least 5 (total) or 1 (maternal or paternal) sequencing reads were assessed for methylation levels and genome-wide tracks were subsequently generated for visualization. The R package DSS v2.14.0 was used to identify DMRs from replicate-merged allele-specific epiblast and EPC CpG report files with smoothing enabled and default parameters. CpGs that showed greater than 10% differential methylation levels between parental alleles with an associated p-value <0.001 were grouped into DMRs (**Sup. table 2**). DMR calls from epiblast and EPC samples were merged using bedtools merge. DNAme levels were then re-calculated from oocytes, sperm, as well as epiblast and EPC samples using bedops bedmap v2.4.39 (Neph et al., 2012). A final list of DMRs wherein all samples are covered by at least 2 CpGs and showed a parental difference of ≥50% DNAme was created (**Sup. Fig. 1b & Sup. table 2**).

For each imprinted gene identified in the allele-specific RNAseq analysis, bedtools Closest v2.27.0 was used to determine the nearest DMR to genic transcription start site. DMRs located within 5kb upstream of an imprinted gene’s TSS were called as promoter DMRs. The *Sall1* DMR was determined by manual curation of an upstream ERVK promoter element. The *Magel2* DMR, which is shared with the *Snrpn* gene, was determined based on conservation with mouse and human DMRs as *Magel2* is incorrectly assembled in the rn6 reference.

### CUT&RUN sequencing analysis

Sequencing reads were trimmed using Trimmomatic (v0.32) and the following parameters: SLIDINGWINDOW:3:10 MINLEN:20 ILLUMINACLIP:TruSeq2_SE.fa:2:30:10. Reads that survived trimming were aligned to genome build mm10 or rn6 using Bowtie2 (v2.2.3) and PCR duplicate reads were flagged using Picard MarkDuplicates (v1.92). Library quality was assessed using samtools flagstat (v1.1). Uniquely aligned, non-PCR-duplicate reads were used for bigwig creation using deeptools bamCoverage (v3.3.0) and parameters: --binSize 1,50,200 -- smoothLength 0,100,2000 minMappingQuality 10 --normalizeUsing CPM –ignoreDuplicates -- ignoreForNormalization chrX chrM chrY --outFileFormat bigwig.

### Comparison of mouse and rat oocyte CUT&RUN data

Rat oocyte H3K4me3, H3K27me3 and H3K36me3 CUT&RUN Spearman correlation metrics were calculated using RPKM levels over 10kb genomics bins and the Morpheus Similarity Matrix tool. Bins with at least 10 CpGs covered by at least 5 sequencing read alignments separated by over 1 read length (n=246,622/285,920, 86%) were considered. Conservation of epigenetic modification profiles in oocytes was measured by Spearman correlation of RPKM and DNAme levels over 1kb rat-mouse syntenic regions generated previously (39) using the Morpheus Similarity Matrix function. Differential enrichment of H3K27me and H3K36me3 levels as well as differential DNAme levels were calculated by subtracting the mouse signal from the rat and plotted as a 2D scatterplot using matplotlib pyplot. A contour plot illustrating the density of data points was generated using seaborn kdeplot.

Rat GVO H3K4me3, H3K27me3, H3K36me3, DNAme (39) and RNAseq (39) data was compared to mouse MII oocyte H3K4me3 (62), H3K27me3 (103), H3K36me3 (58), DNAme (42) and RNAseq (62) over transcribed and repressed genes in oocytes. Gene expression levels in the oocyte were calculated using the rat-mouse homologous gene annotation described above. For each species, genes were categorized as expressed (RPKM ≥1) or not (RPKM<1), and heatmaps were generated over both groups using deeptools ComputeMatrix and PlotHeatmap functions for both species.

To directly compare rat, mouse and human oocyte H3K4me3 peak sizes, human and mouse oocyte H3K4me3 CUT&RUN data were downloaded from (64) as well as rat (60) and reprocessed using the same parameters. SICER2 (110) was used with default parameters to call peaks and peaks within 3kb were grouped together. The distribution of H3K4me3 peak sizes was plotted using VisRseq and peak call files were converted to bigBed for visualization in the UCSC Genome Browser (111).

### Software used

Bedops (v2.4.39) (112)

Bedtools (v2.22.1) (113)

Bismark (v0.16.3) (114)

Bowtie2 (2.2.3) (115)

Deeptools2 (v3.3.0) (116)

DSS (v2.14.0) (37, 38)

Limma (v3.50.1) (108)

MEA (v1.0) (26)

Morpheus (Broad Institute)

PicardTools(v1.92) (117)

R base (v4.1.2) (109)

Samtools (v1.1) (118)

Seaborn (119)

SICER2 (110)

STAR (v2.4.0.i) (120)

Stringtie (v2.1.4) (105)

Trimmomatic (v0.32) (104)

UCSC Genome Browser (111)

UCSC Track Hubs (121)

VisRseq (v0.9.12) (122)

## DECLARATIONS

## Ethics approval

All experiments were performed in accordance with the animal care and use committee guidelines of the National Institute for Physiological Sciences and Tokyo University of Agriculture.

## Consent for publication

Not applicable

## Availability of data and materials

Sequencing datasets generated in this study have been deposited in GEO under the accession numbers:

**GSE185573** Rat oocyte epigenomes

**GSE185574** Rat and mouse imprintomes (allele-specific transcriptomes)

**GSE186492** Rat and mouse imprintomes (allele-specific DNA methylomes)

See Supplemental table 2 for the full list of data analyzed for this study.

All datasets analyzed in this study can be visualized on the UCSC Genome Browser: https://genome.ucsc.edu/s/JRA/the_rat_imprintome_share

Publicly available software and VisRseq (v0.9.12) were used for data analysis and plot generation. Custom ad-hoc bash and python scripts were also generated for data filtering and plotting and have been deposited in GitHub and are available at: https://github.com/julienrichardalbert/the_rat_imprintome/releases/tag/v0.1 and https://github.com/kdkorthauer/rat_imprintome/releases/tag/initial-submission

## Authors’ contributions

T.K, M.O., M.H., S.T and H.K. prepared the rat embryos. S.K. and H.K. prepared the mouse embryos. S.K., T.T., A.M., and H.K. generated the RNAseq, WGBS and PBAT sequencing libraries. A.I. generated CUT&RUN libraries. F.M. generated tPBAT libraries. A.M.S. performed the human literature search. K.Korthauer informed statistical analyses. M.C.L., K.Kurimoto., M.C.V.G. and H.K. supported the project. J.R.A. processed the data, performed analyses, generated the figures and wrote the manuscript with M.C.L.

## Acknowledgements

We are grateful to Louis Lefebvre (University of British Columbia) for critical reading of the manuscript. We acknowledge Dr. Keisuke Tanaka (Tokyo University of Agriculture) for library preparation and quality control and Dr. Akihiko Sakashita (KEIO University) for help preparing embryos. This research was enabled in part by support provided by Compute Canada (www.computecanada.ca) and GenAP (genap.ca).

## SUPPLEMENTAL TABLES

**Supplemental table 1** Data displayed in figure panels.

**Supplemental table 2** Rat differentially methylated regions.

**Supplemental table 3** Datasets generated and mined in this study.

## ADDITIONAL FILES

**Additional Data 1** Allele specific RNAseq

**Additional Data 2** Allele agnostic RNAseq

**Additional Data 3** Syntenic 1kb regions oocyte epigenetic profiling

**Additional Data 4** Rat and mouse oocyte 1kb bin DNAme levels

**Additional Data 5** Rat 10kb bins epigenetic profiling

