## Supplemental figures and legends for "Conservation and divergence of canonical and non-canonical imprinting in murids"

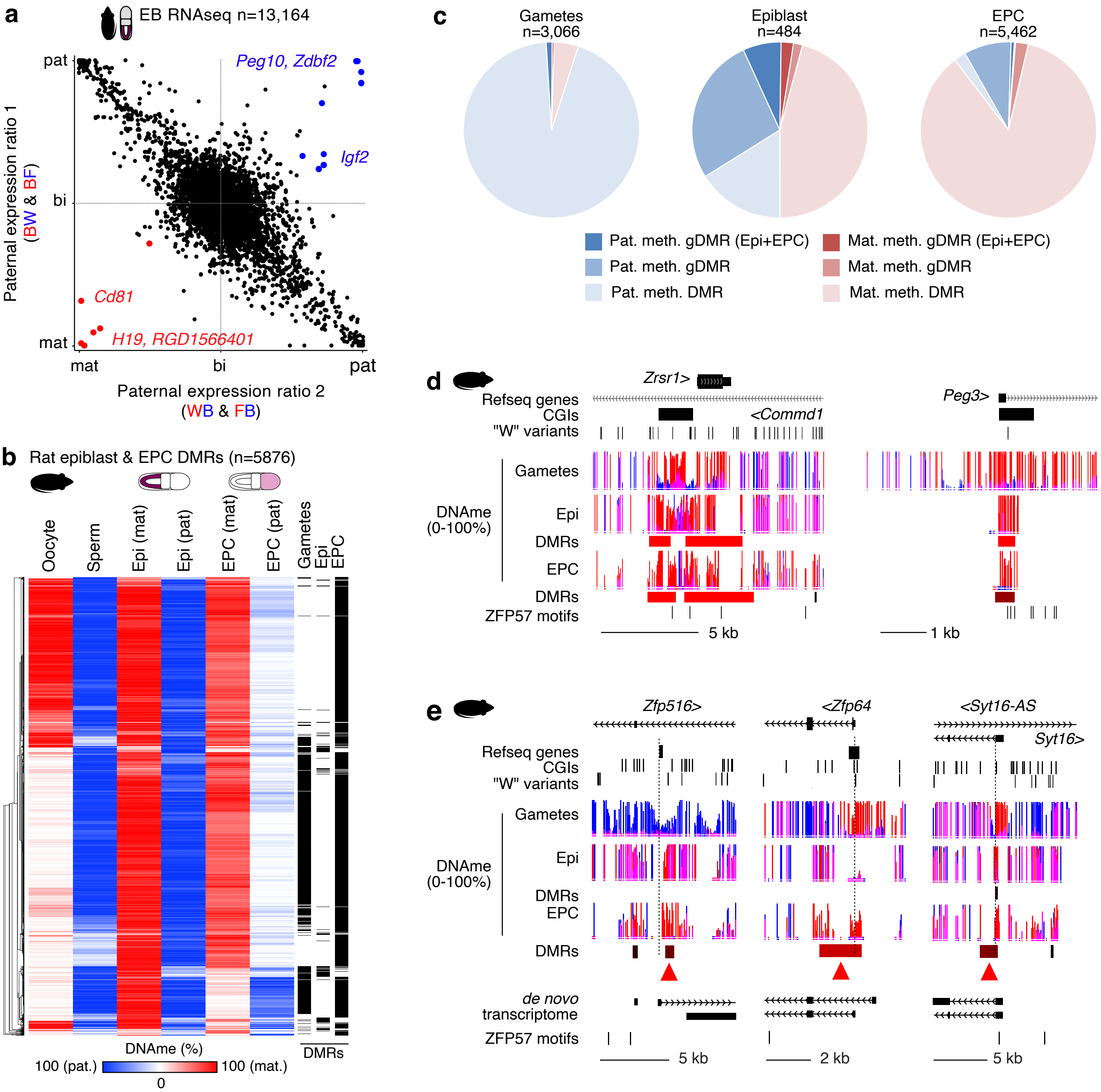


**Supplemental Figure 1. Agnostic identification of rat differentially methylated regions (DMRs) in rat epiblasts and EPCs.** **a** Scatterplot displaying paternal expression ratios of transcripts in rat epiblast (Epi) cells as in **Fig. 1b**. The paternal expression ratio was averaged over 11 samples generated from two distinct reciprocal crosses, and expressed transcripts (RPKM ≥1) with sufficient allelic coverage (RPM ≥0.5) in at least 6 samples were reported (n=13,164). Genes showing parent-of-origin imprinted gene expression (Student's T-test, Bonferroni-adjusted p-val <0.05) are coloured red (maternally expressed) or blue (paternally expressed). **b** Heatmap showing gamete and parent-specific Epi and ectoplacental cone (EPC) DNAme levels over DMRs (n=5,877). Hierarchical clustering was performed based on DNAme levels in all samples. **c** Pie charts showing the number of paternally and maternally methylated DMRs in rat gametes, epiblasts and EPCs. DMRs are further classified by their imprinting status (>50% parental genome DNAme level difference) in gametes (gametic DMR, gDMR) and all samples (gDMR + Epi + EPC). **d-e** Rat genome browser screenshots of the *Zrsr1/Commd1, Peg3, Zfp516, Zfp64* and *Syt16-AS* loci showing maternally methylated differentially methylated regions (DMRs) in gametes, Epi and/or EPC. The genomic position of known Refseq genes, CpG islands (CGIs) as well as rat parental strain SNVs and INDELs are included. Note the single variant in the *Peg3* CGI promoter DMR was sufficient to correctly call maternal-specific methylation. DNA methylation (DNAme) levels are represented as bar charts for each CpG dinucleotide with at least 5 allele-agnostic (gametes) or 1 allele-specific (Epi/EPC) sequencing read alignment. Maternal and paternal DNAme bar charts are coloured red and blue, respectively, and CpGs with maternal and paternal coverage are in pink. The DMR tracks are coloured as a function of statistical significance and differential parent-allele DNAme levels.

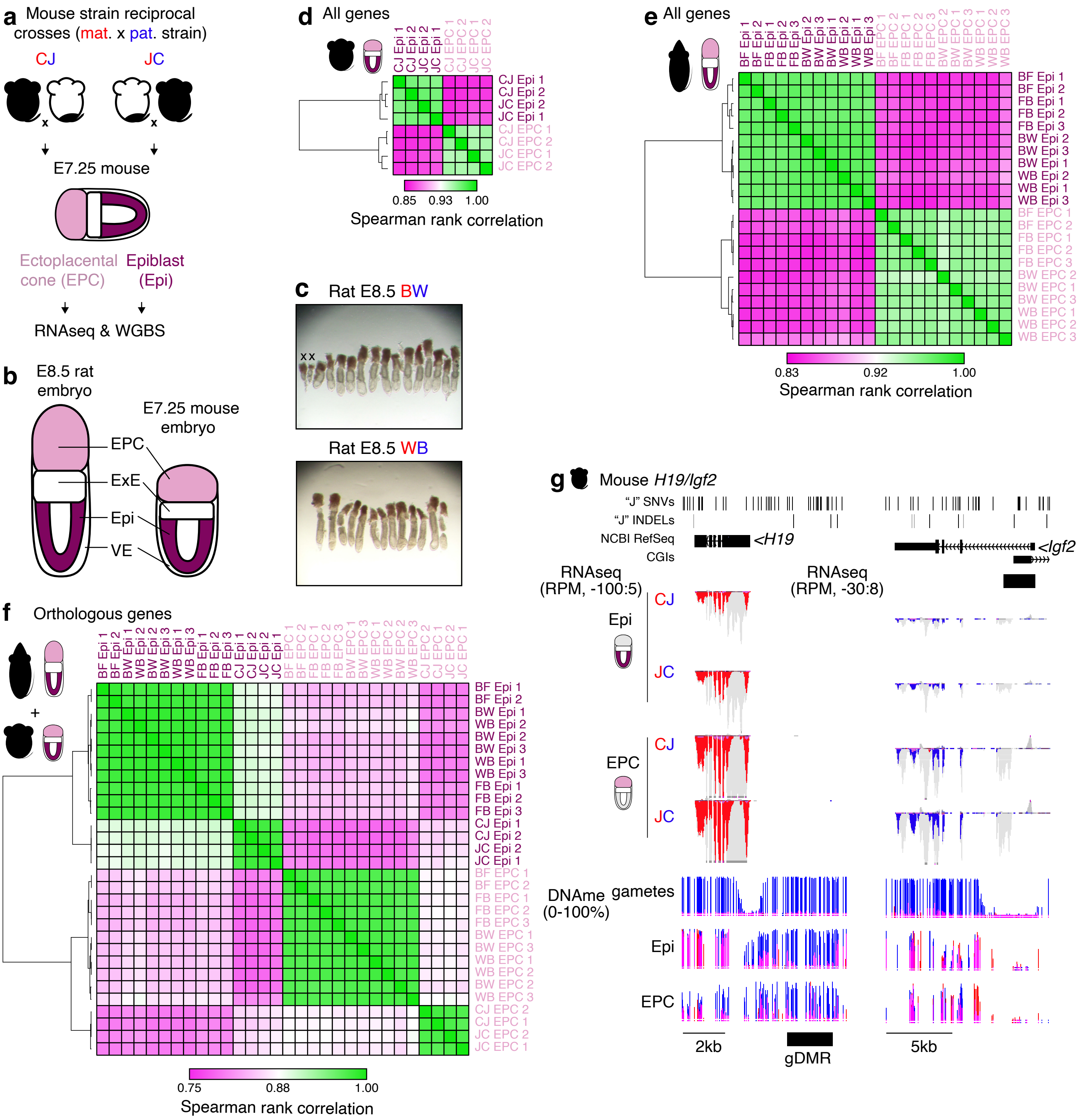


**Supplemental Figure 2. RNAseq analysis of rat and mouse embryos.**
**a** Experimental design of mouse matings and tissue collection. Reciprocal crosses of distinct mouse strains C57BL/6N, "C" and JF1/Ms, "J" were performed and cells from the E7.25 epiblast and EPC were collected. Strand-specific RNAseq and WGBS was performed on all samples in duplicate. **b** Schema comparing rat and mouse embryos at Carnegie stage 7, corresponding to E8.5 in and and E7.25 in mouse. Sequenced tissues are highlighted. EPC: ectoplacental cone, ExE: extraembryonic ectoderm, Epi: epiblast, VE: visceral endoderm. **c** Rat embryo imaging. Note BN/NCrlCrlj, "B", females produce more inviable offspring relative to WKY/NCrlCrlj, "W", as indicated by crosses. **d** Correlogram showing clustering of mouse Epi and EPC transcript expression. Samples were correlated by Spearman rank correlation of transcript expression values (n=106,520) agnostic to allelic information. **e** Correlogram showing clustering of rat epiblast (Epi) and ectoplacental cone (EPC) samples. Correlogram and scale as in **d** over rat transcripts (n=69,193). **f** Correlogram showing clustering of rat and mouse samples. Samples were correlated by Spearman rank correlation of rat and mouse orthologous gene expression values (n=16,769) agnostic to allelic information. **g** Mouse genome browser screenshots of the *H19* and *Igf2* loci. The genomic position of known Refseq genes, CpG islands (CGIs) as well as mouse parental strain SNVs and INDELs are included. For each cross, biological duplicate samples were merged and the mean expression level is displayed in reads per million (RPM). Read alignments (grey) are highlighted if they align to the maternal (red) or paternal (blue) alleles. CpG DNAme levels are reported as bar graphs, and the location of each CpG with sufficient read coverage (5 for allele-agnostic and 1 for maternal or paternal alleles) is indicated under each bar graph. Maternal and paternal DNAme bar charts are coloured red and blue, respectively, and CpGs with maternal and paternal coverage are in pink. The location of known mouse gametic DMRs (gDMRs) (101) and scale bars are included.

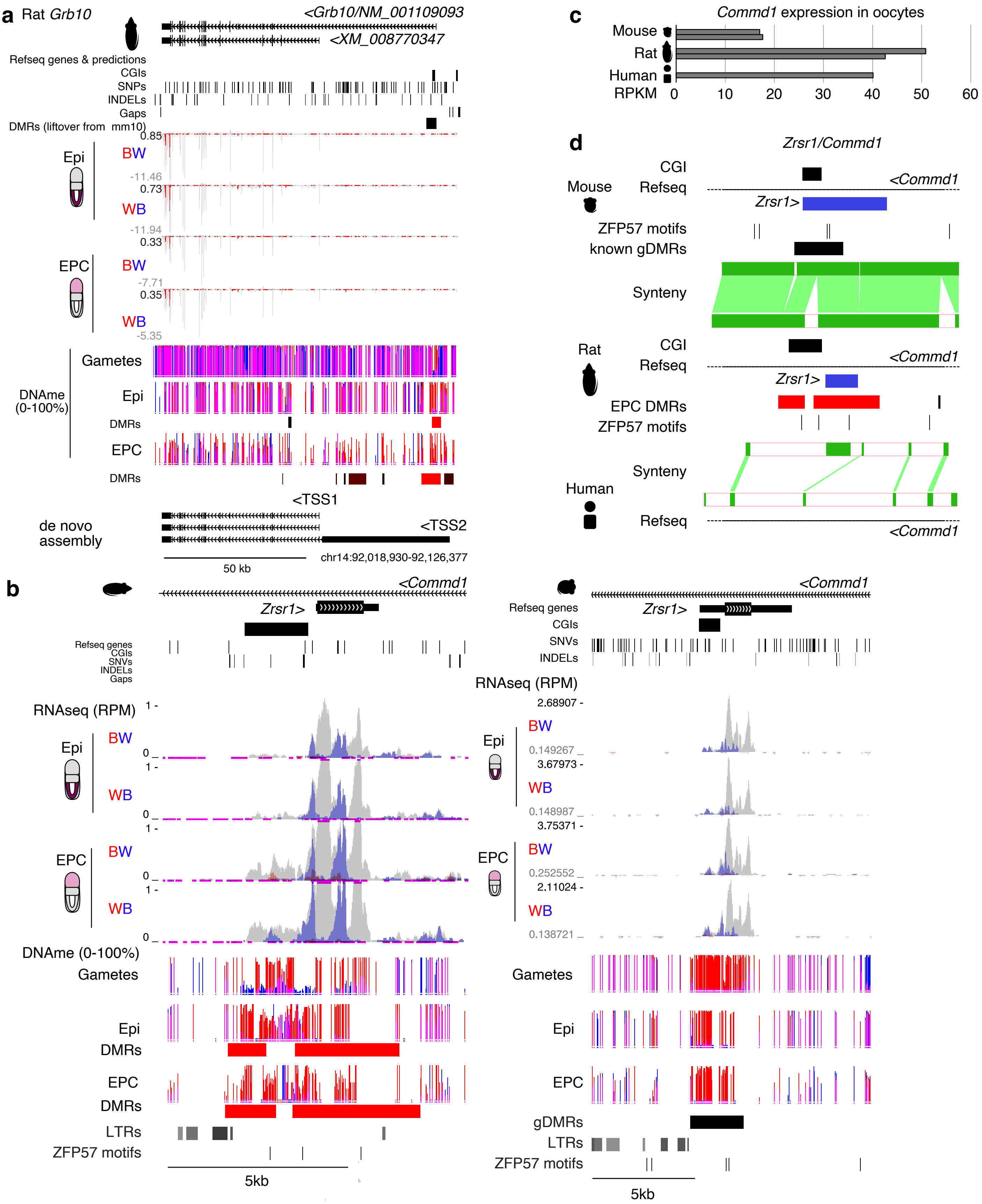


**Supplemental Figure 3. Maternal expression of rat *Grb10* and nearby DMR.**
**a** Rat genome browser screenshot of the *Grb10* locus showing maternal specific expression and a maternally methylated differentially methylated region (DMR) in gametes, epiblast (Epi) and ectoplacental cone (EPC). Tracks are presented as in **Sup. Fig. 2g**. The location of known mouse gametic DMRs (gDMRs), *de novo* transcriptome assembly and scale bars are included. **b** Rat and mouse genome browser screenshots of the *Zrsr1/Commd1* locus showing paternal specific expression of *Zrsr1* and a maternally methylated CGI promoter in gametes, Epi and EPC. Data tracks as in **a**. **c** *Commd1* expression levels in mouse, rat and human oocytes are shown in RPKM. **d** Ensembl Region Comparison screenshot of intron 1 of *Commd1* in mouse, rat and human, including the *Zrsr1* gene (blue). Sequence synteny is illustrated as green bars. All other tracks are as shown in **b**.

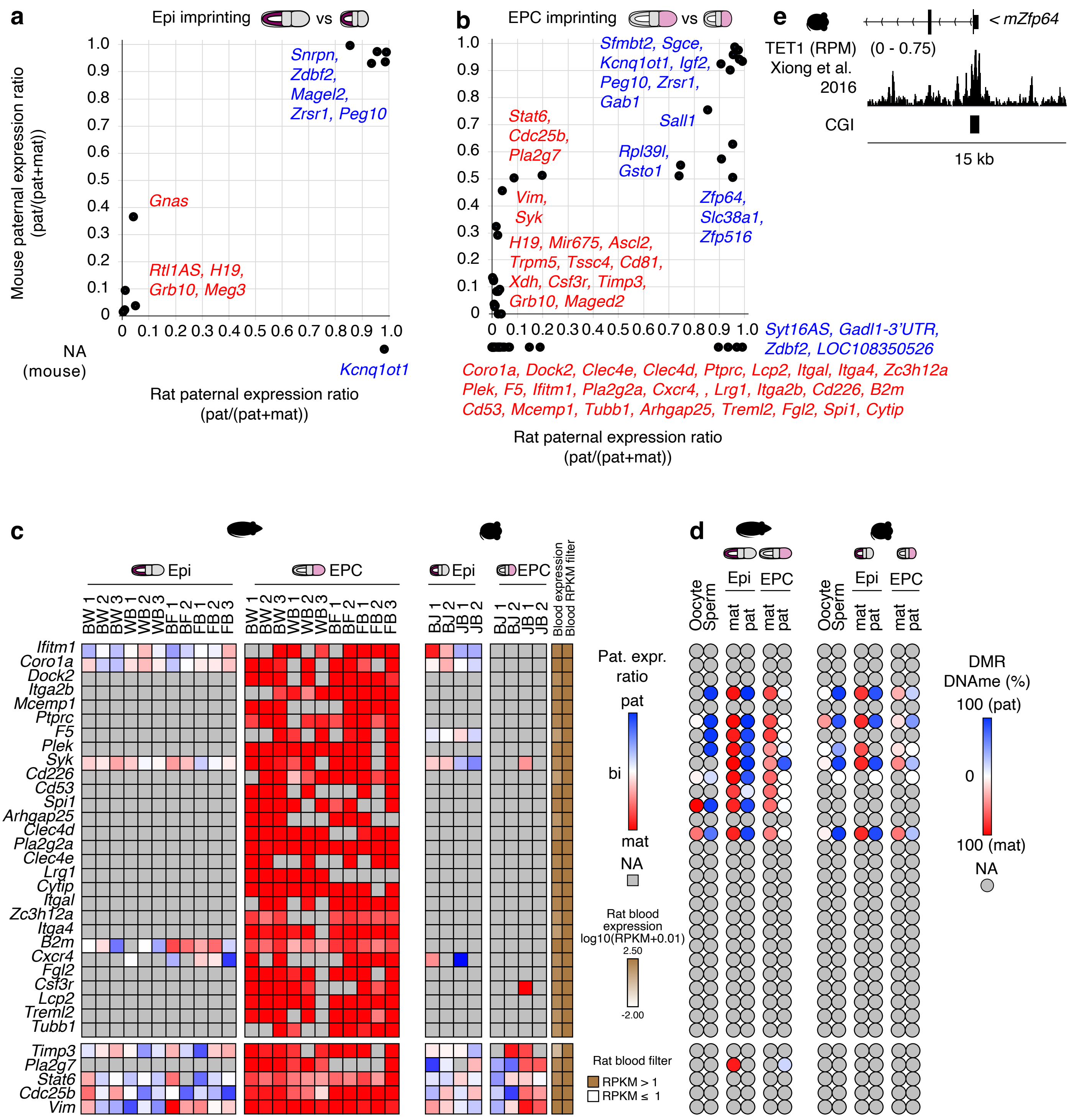


**Supplemental Figure 4. Conserved and rat-specific imprinted gene expression.**
**a** 2D scatterplot showing parental expression ratios in rat versus mouse epiblast (Epi) cells. The average paternal expression ratio for each gene is shown. Expressed (RPKM ≥1) rat genes with sufficient allele-specific read coverage (allelic RPM ≥0.5) and show statistically significant bias in monoallelic gene expression (Bonferroni-adjusted p-value <0.05, Student's t-test) are shown. For genes with insufficient information to ascertain mouse expression ratios, the rat paternal expression ratio is shown below the x-axis. **b** 2D scatterplot showing parental expression ratios in rat versus mouse ectoplacental cone (EPC) cells. **c** Heatmap of paternal expression ratios in rat and mouse Epi and EPC cells for genes that show maternal expression uniquely in EPC and are normally expressed (RPKM>1) in adult rat blood. None of the reported genes are known to be imprinted in human. **d** Heatmap of allele-specific DNAme levels over DMRs associated with putative imprinted genes in **c**. **e** Mouse genome browser screenshot of the mouse *Zfp64* CGI promoter showing enrichment of the TET1 enzyme in reads per million (RPM). TET1 ChIP-seq downloaded from (102).

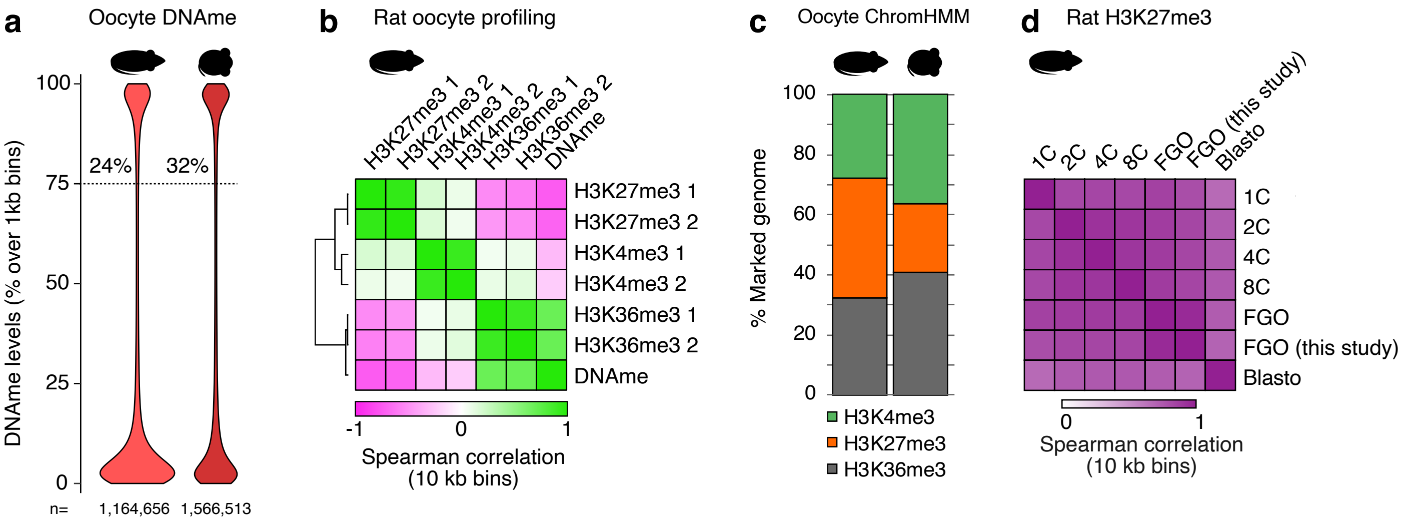


**Supplemental Figure 5. Epigenome profiling of rat oocytes by CUT&RUN.**
**a** Violin plot showing the genome-wide distribution of DNAme levels (over 1kb bins) in rat and mouse oocytes. The percent of 1kb bins with >75% DNAme is indicated. **b** Correlogram showing genome-wide Spearman correlation metrics between H3K4me3, H3K27me3 and H3K36me3 (CUT&RUN) and DNAme levels over 10kb bins in rat oocytes. **c** The percentage of the genome marked by H3K4me3, H3K27me3 and H3K36me3 in rat and mouse oocytes is shown. **d** Genome-wide Spearman correlation of H3K27me3 levels over 10kb bins in rat oocytes and preimplantation embryos from (60) are shown.

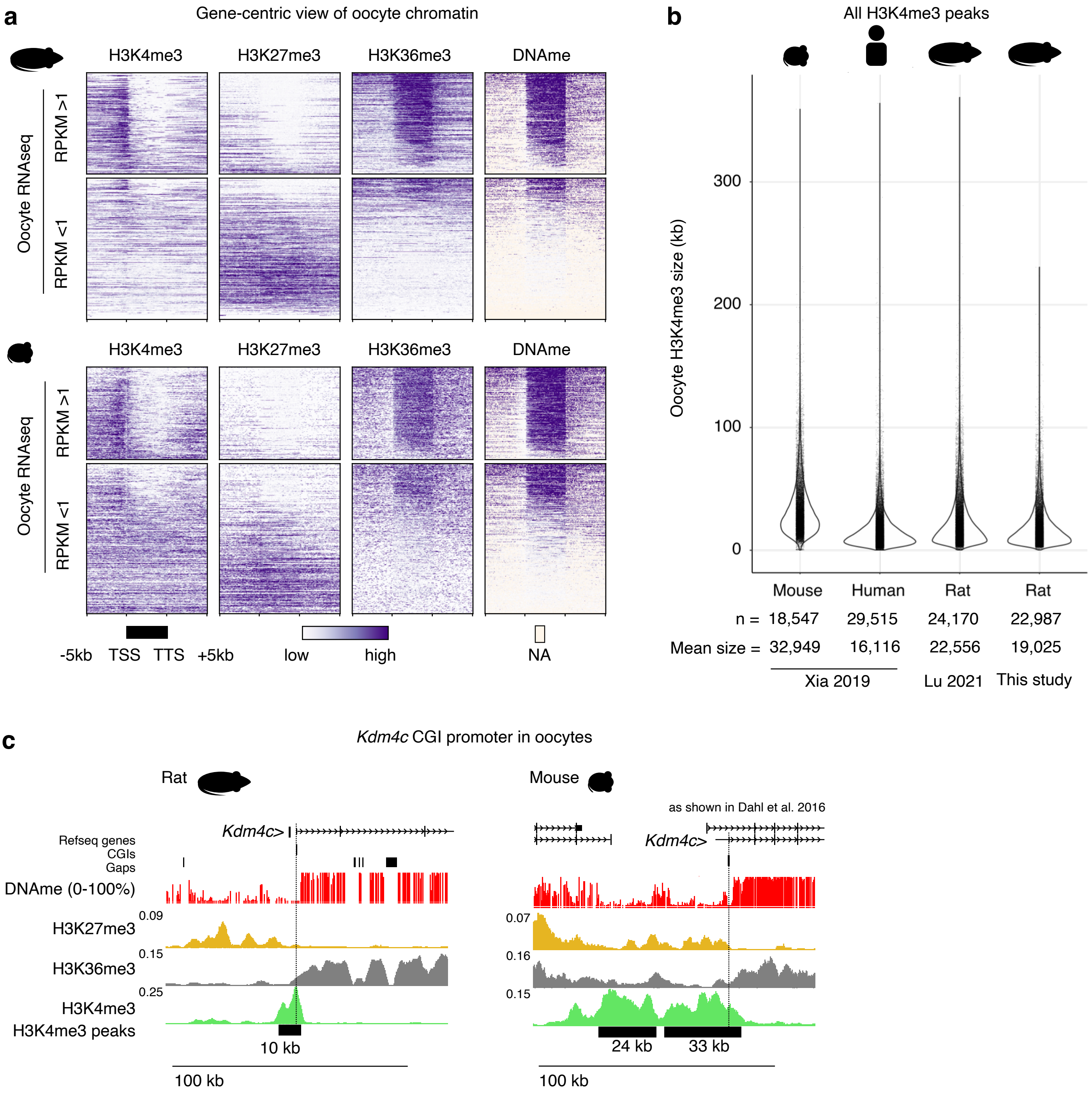


**Supplemental Figure 6. Oocyte domain sizes in rat, mouse and human oocytes.**
**a** Gene-centric heatmaps of mouse and rat oocyte epigenomes. Genes are grouped by their expression level (RPKM ≥1 and RPKM <1) in oocytes. **b** Violin plot showing the distribution of mouse, human and rat oocyte H3K4me3 domain sizes in kilobases (kb). The number of peaks and average size is indicated below each violin plot. Mouse and human data from (64). Rat data from (60) and this study. **c** Genome browser screenshots of the *Kdm4c* CGI promoter in rat and mouse oocytes as previously shown (61). The transcription start site of *Kdm4c* is indicated by a dashed line. Oocyte ChIP or CUT&RUN data are shown in counts per million (CPM). The approximate size of H3K4me3 domains are indicated.

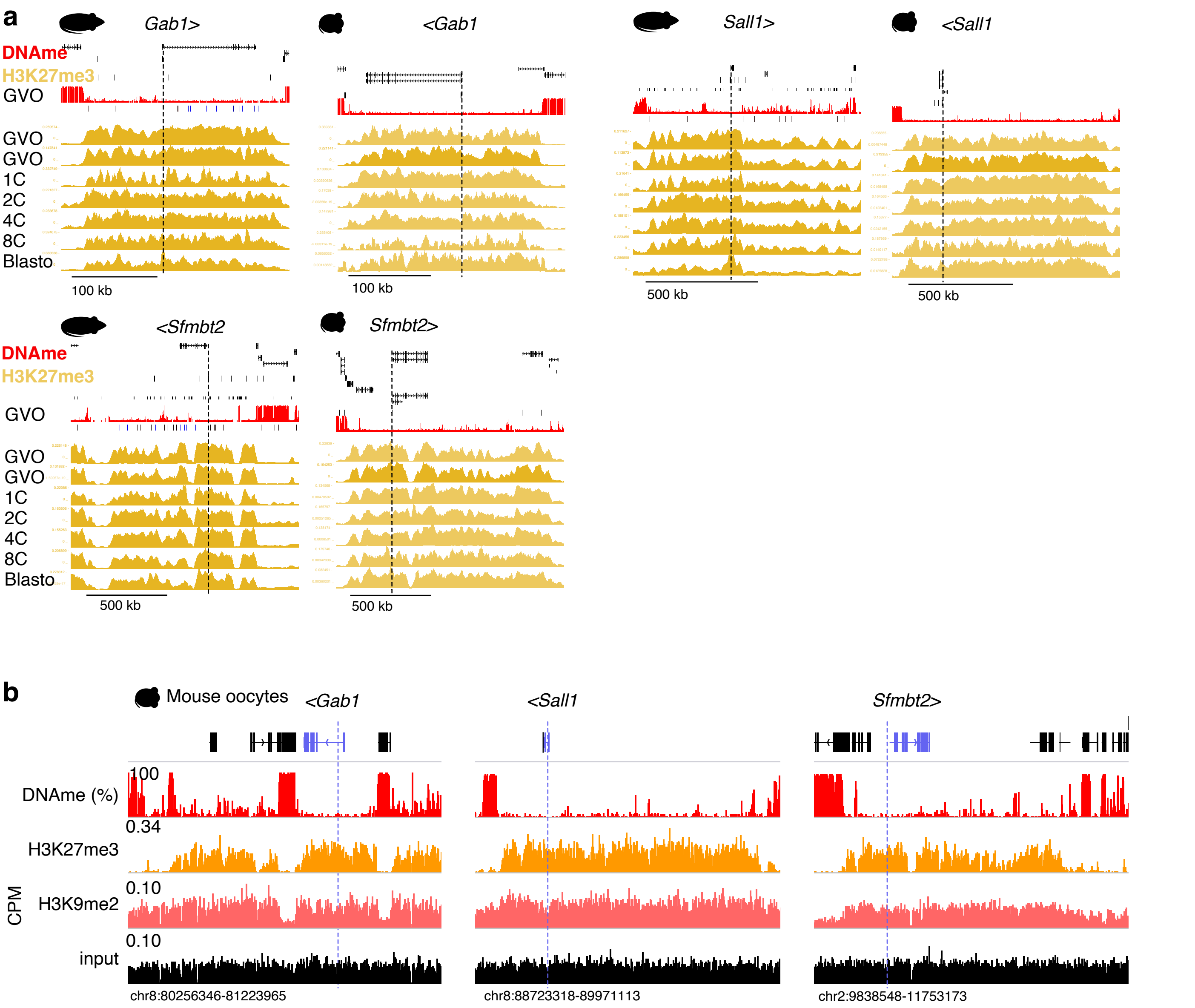


**Supplemental Figure 7. Establishment and maintenance of conserved non-canonical H3K27me3 imprints in rat and mouse preimplantation embryos.**

**a** Genome browser screenshots of conserved non-canonically imprinted genes in rat and mouse *Gab1, Sall1* and *Sfmbt2* .Tracks are presented in the following order: Refseq genes, CpG islands (CGIs), gaps in the reference genome (rat only), oocyte DNA methylation levels (red, 0-100%), oocyte H3K27me3 CUT&RUN (this study), as well as oocyte H3K27me3 CUT&RUN, 1C, 2C, 4C, 8C, blastocyst (blasto) in rat from (60) and mouse from (103) (orange, counts per million). The approximate location of promoters of non-canonically imprinted, paternally expressed genes are indicated by dotted lines. **b** Genome browser screenshots of *Gab1*, *Sall1* and *Sfmbt2* in mouse oocytes. Scales are in Counts Per Million (CPM). Dotted blue lines indicate maternally methylated DMRs in mouse EPCs.

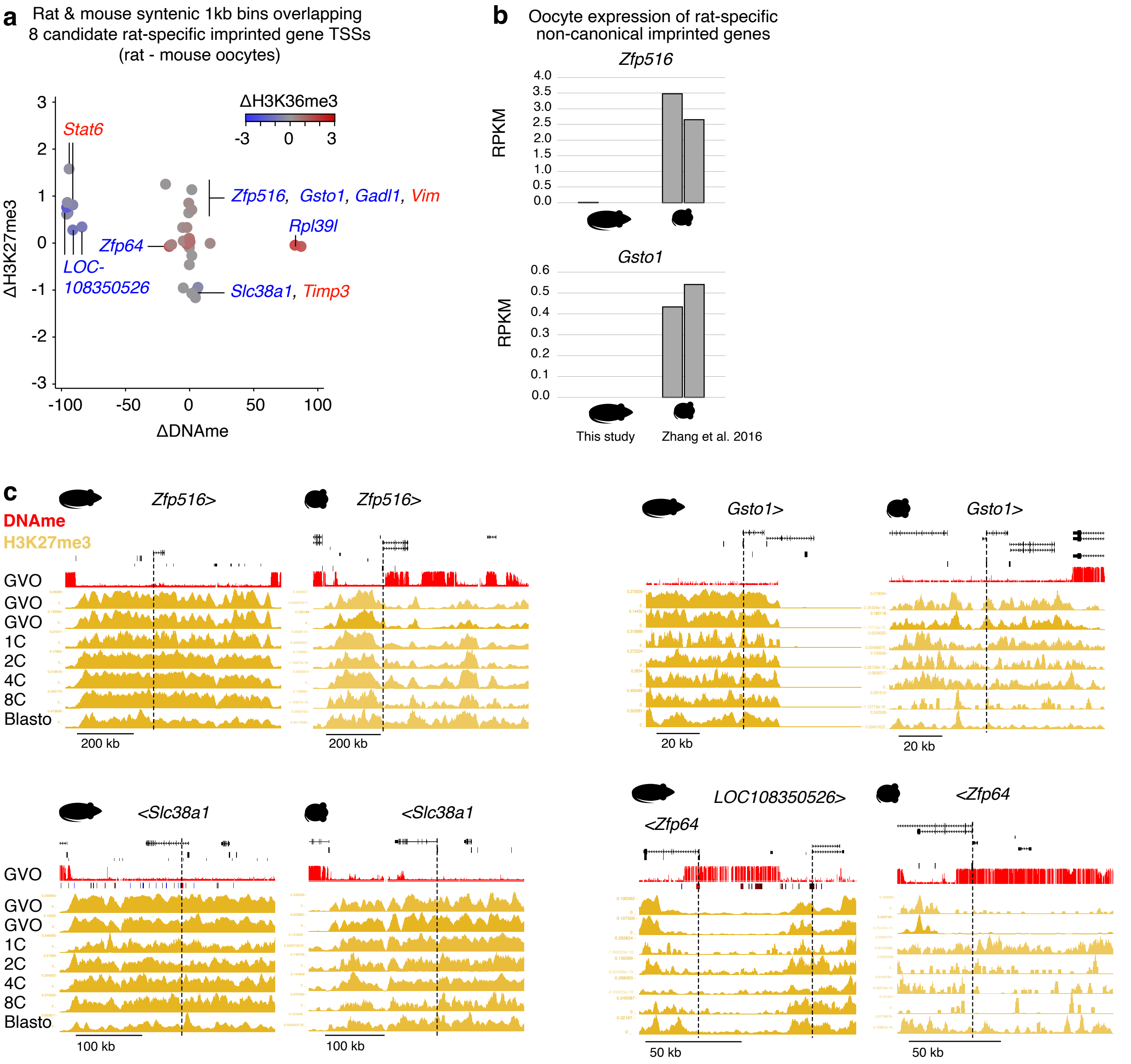


**Supplemental Figure 8. Epigenetic and expression profiling of novel rat-specific non-canonical imprinted genes in rat and mouse oocytes and preimplantation embryos.**

**a** Scatterplot showing differences in rat and mouse oocyte DNAme, H3K27me3 and H3K36me3 levels over syntenic 1kb bins overlapping and adjacent to the DMR or TSS of non-canonical rat-specific imprinted genes. **b** Bar charts showing expression levels of rat-specific imprinted genes *Zfp516* and *Gsto1* in rat and mouse germinal vesicle oocytes (GVOs). **c** Genome browser screenshots of rat-specific non-canonically imprinted genes *Zfp516*, *LOC108350526*, *Slc38a1* and *Gsto1.* Tracks are presented in the following order: Refseq genes, CpG islands (CGIs), gaps in the reference genome (rat only), oocyte DNA methylation levels (red, 0-100%), oocyte H3K27me3 CUT&RUN (this study), as well as oocyte H3K27me3 CUT&RUN, 1C, 2C, 4C, 8C, blastocyst (blasto) in rat from (60) and mouse from (103) (orange, counts per million). The approximate location of promoters of non-canonically imprinted, paternally expressed genes are indicated by dotted lines.

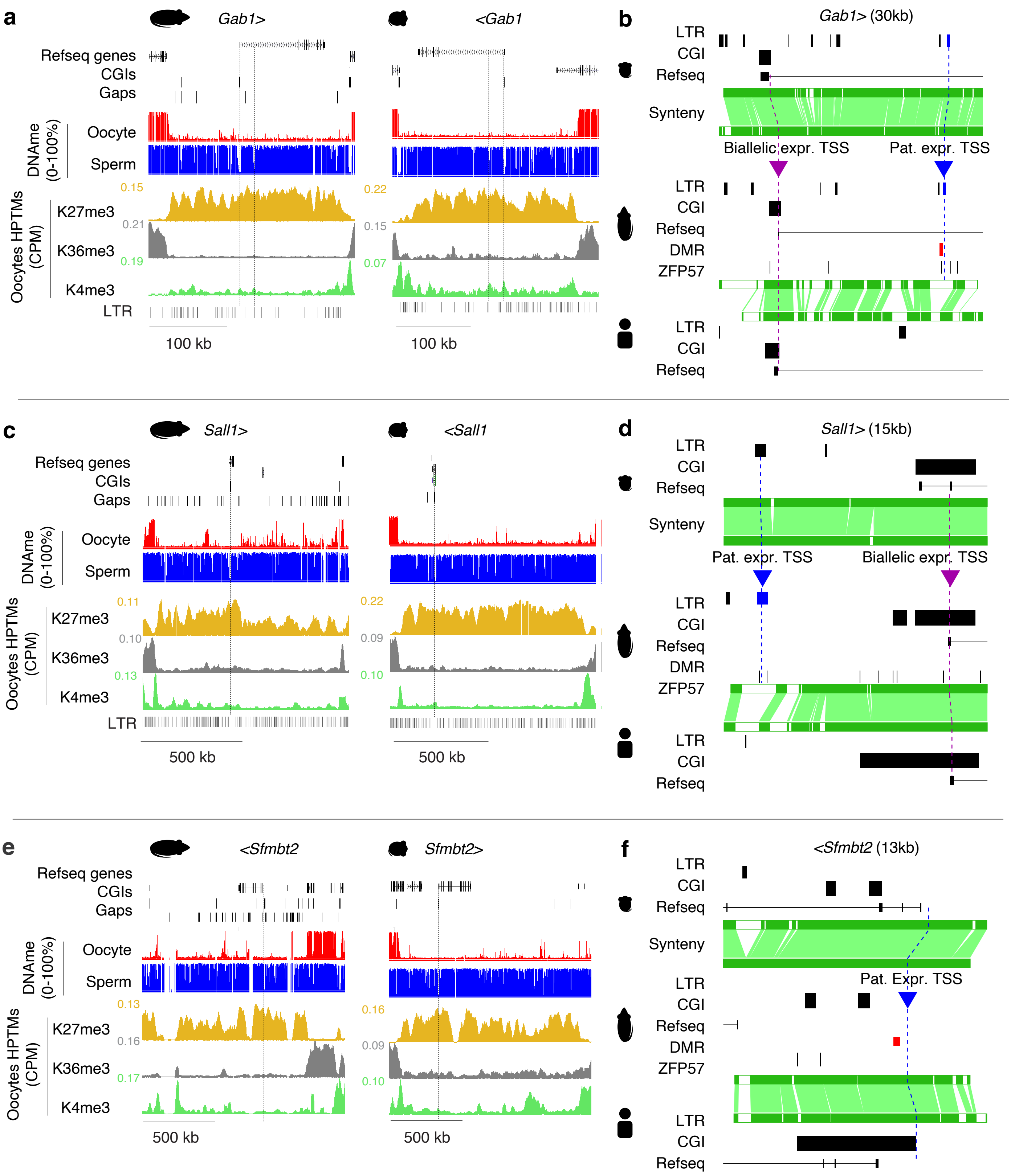


**Supplemental Figure 9. Oocyte epigenome profiling of conserved non-canonical imprinted genes *Gab1*, *Sall1* and *Sfmbt2* and syntenic promoter analysis.**
**a** Rat and mouse genome browser screenshots of the *Gab1* CGI promoter. The position of the canonical annotated transcription start site (TSS) is indicated by a dashed line. The location of the alternative paternally expressed ERVK promoter is indicated by an intronic dashed line. Oocyte CUT&RUN (rat) and ChIP (mouse) data are shown in counts per million (CPM). DNAme levels in oocytes (red) and spermatozoa (blue) are included. CGIs, gaps in the rat genome, and long terminal repeat (LTR) elements are indicated. **b** Ensembl Region Comparison screenshot of the *Gab1* canonical CGI and alternate ERVK promoters in rat (rn6), mouse (mm10) and human (hg19). Syntenic sequences shown in green. The locations of mouse, rat and human Refseq genes, LTRs and CGIs are shown. Rat EPC DMRs and ZFP57 binding motifs are shown. Canonical (biallelic, purple) and alternate (paternal-specific, blue) promoters are indicated by triangles and their conservation across species is shown by a dashed line. **c-d** Oocyte epigenome profiling and syntenic sequence analysis of the *Sall1* locus. **e-f** Oocyte epigenome profiling and syntenic sequence analysis of the *Sfmbt2* locus.
